# Learning the structure of the world: The adaptive nature of state-space and action representations in multi-stage decision-making

**DOI:** 10.1101/211664

**Authors:** Amir Dezfouli, Bernard W. Balleine

## Abstract

State-space and action representations form the building blocks of decision-making processes in the brain; states map external cues to the current situation of the agent whereas actions provide the set of motor commands from which the agent can choose to achieve specific goals. Although these factors differ across environments, it is not currently known whether or how accurately state and action representations are acquired by the agent because previous experiments have typically provided this information a priori through instruction or pre-training. Here we show that, in the absence of such a priori knowledge, state and action representations adapt to reflect the structure of the world. We used a sequential decision-making task in rats in which they were required to pass through multiple states before reaching the goal, and for which the number of states and how they map onto external cues were not known a priori. We found that, early in training, animals selected actions as if the task was not sequential and outcomes were the immediate consequence of the most proximal action. During the course of training, however, rats recovered the true structure of the environment and made decisions based on the expanded state-space, reflecting the multiple stages of the task. We found a similar pattern with actions; early in training animals only considered the execution of single actions whereas, after training, they created useful action sequences that expanded the set of available actions. We conclude that the profile of choices shows a gradual shift from simple representations of actions and states to more complex structures compatible with the structure of the world.

## 1 Introduction

In sequential decision-making tasks an agent makes a series of choices and passes through several states before earning reward. Such tasks involve planning and learning the consequences of specific actions. Within the lab, however, important aspects of the environment, such as the available actions, the number of states of the environment and how they map on to external cues, i.e., the state-space of the task, are typically given to subjects in advance of the task either through instructions or pre-training. Such a priori information is, however, not typically made available to a decision-maker in natural environments, and the decision-making process must, therefore, involve (i) learning the correct state-space of the environment and (ii) acquiring new actions that are useful for earning reward in the task.

Learning the state-space of the task is, therefore, crucial in allowing the agent to navigate within the environment, and provides building blocks for various forms of reinforcement-learning algorithms in the brain (Sutton and Barto, 1998; Ito and Doya, 2011). This process involves considering different events and cues that occur after taking each action, and integrating them in order to recover how many states the task has and how they are related to external cues. Recent theoretical work provides a basis for this process in the context of classical conditioning and suggests that the underlying states used to represent the environment are dynamic; that animals are able to infer and learn new states of the environment based on their observations (Gershman et al., 2010; Redish et al., 2007). However, at present, there is no direct evidence for such adaptive state-space representations in decision-making situations.

Actions are the other building block for reinforcement-learning algorithms, and refer, for example, to different motor commands that an agent can use to influence the state of the environment. Efficient decision-making relies on using actions at the appropriate scale; engaging decision-making at too fine-grained a level of motor movement will overwhelm this process with choice points. Nevertheless, evidence suggests that humans and other animals can create new actions in the form of action chunks or action sequences by concatenating simple actions together (Botvinick et al., 2009; Dezfouli and Balleine, 2012; Smith and Graybiel, 2014; Lashley, 1951; Ito and Doya, 2011). Such action sequences, known as ‘temporary extended actions’, can be thought of as new skills that expand the set of available actions and that are treated as single response units. By acquiring new action sequences, the selection process needs to be implemented only once at the initiation point of an action sequence instead of before each individual action and, in this way, adaptive representations of actions contribute to the scalability of the decision-making process.

In the current study, using a sequential decision-making task in rats we sought to investigate whether state-space and action representations adapt to the structure of the world. We used a two-stage decision-making task similar to a two-stage task previously used in human subjects (e.g., Daw et al., 2011; Dezfouli and Balleine, 2013), and show, without any explicit instructions about the structure of the task (which obviously cannot be provided to rats), that early in training, the rats made decisions based on the assumption that the state-space is simple and the environment is composed of a single stage whereas, later in training, they learned the true state-space reflecting the multi-stage structure of the environment and made decisions accordingly. We were then able to show that, concurrently with the expansion of the state-space, the set of actions also expanded and action sequences were added to the set of actions that the rats executed. The evidence shows, therefore, that decision making is adaptive and depends on acquiring, refining and updating both state-space and action representations over time.

## 2 Results

Rats (n=8; see Material and Methods) received several pre-training sessions, in which they learned to press two levers, ‘R’ and ‘L’ (right and left), to earn food pellets (Figure 1:phase 2) and then to discriminate between two stimuli S1 and S2 such that in the presence of S2 action ‘R’ was rewarded whereas in the presence of S1 action ‘L’ was rewarded (Figure 1:phase 3; Figure A2). Subsequently, the rats received training on a two-stage decision-making task, in which they first made a binary choice at stage 1 (S0), after which they transitioned to one of the stage 2 states, (either S1 or S2), in which again they made another binary choice that could lead to either reward delivery or no-reward (Figure 2a). In each trial, only one of the stage 2 states led to reward; the other state did not lead to reward irrespective of the choice of actions (Figure 2b). During the course of each session, the stage 2 state that earned a reward switched without any signal (with probability 0.14; indicated by ‘reversal’ in Figure 2b).

**Figure 1.**
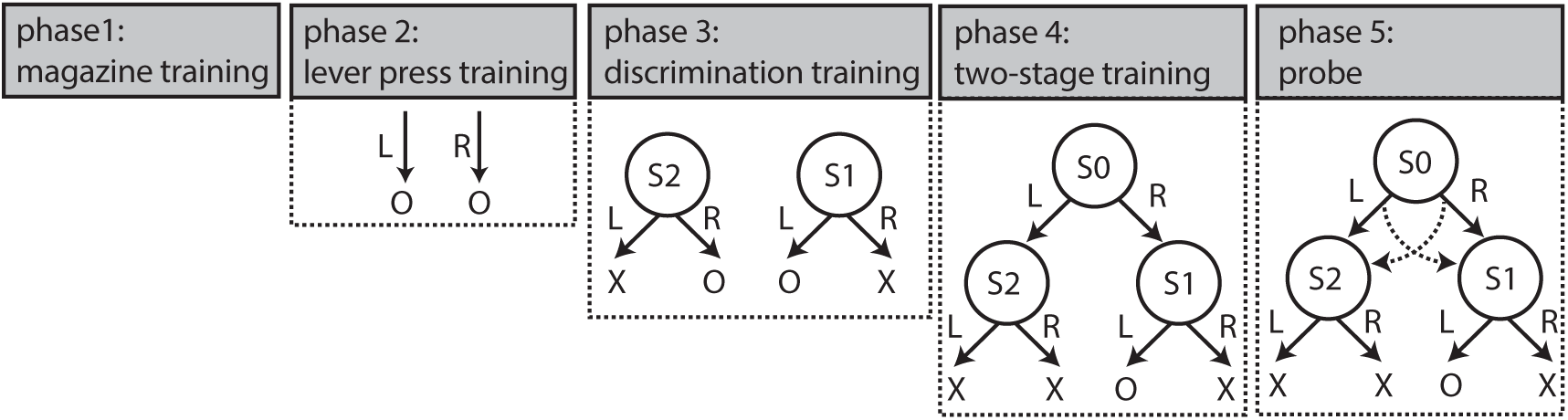
Different phases of the experiment. The experiment started with two magazine training sessions (phase 1), followed by several lever training sessions (phase 2), in which animals learned that pressing each lever (left and right levers corresponding to ‘L’ and ‘R’ in the figure) would delivered a reward (presented by ‘O’ in the figure). The next phase was discrimination training (phase 3), in which animals learned that when stimulus S1 was presented action ‘L’ should be taken to earn a reward, and when S2 was presented action ‘R’ should be taken to earn a reward. S1 and S2 were a constant and blinking house light, respectively. The next phase of the experiment was two-stage training, in which animals were trained on a two-stage decision-making task. This training phase comprised multiple training sessions and, in the middle or at the end of these training sessions, several ‘probe sessions’ were inserted.

**Figure 2.**
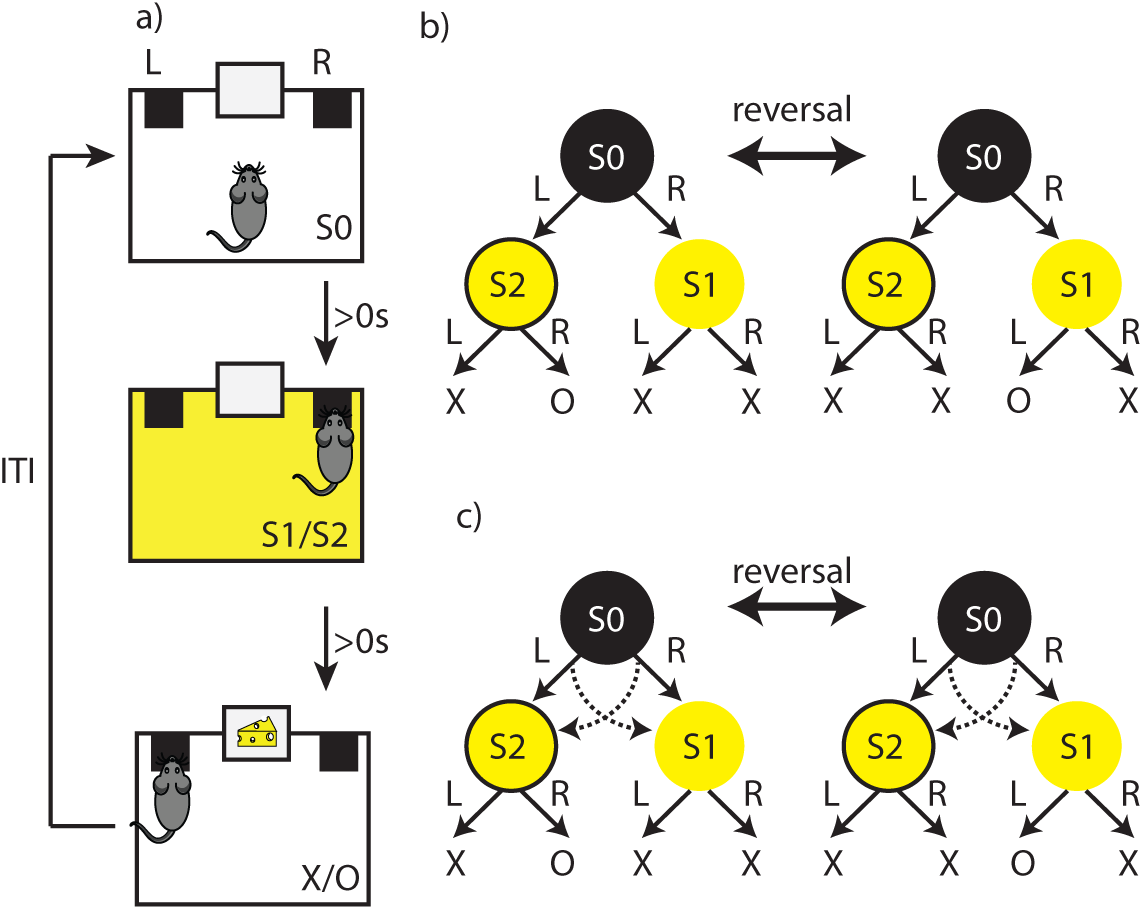
(a) The flow of events in the two-stage task. Trials started in state S0, which was signalled by the absence of the house light. After an action (‘L’ or ‘R’) was taken at stage 1, either constant or blinking house light started (S1 or S2). Next, subjects could take another action at stage 2 (‘L’ or ‘R’), which could lead either to the delivery of the outcome or to no outcome. Actions taken in S0 immediately lead to the presentation of either S1 or S2, and actions taken in S1 or S2 immediately lead to the outcome or no outcome. The inter-trial interval (ITI) was zero in this experiment, but in the experiments reported in the Supplementary Materials, it was greater than zero, as detailed in Supplementary Materials. (b) The structure of the task. Stage 1 actions in S0 led to the stage 2 stimuli (S1/S2) in a deterministic manner. The rewarding stage 2 state changed with a probability of 0.14 after earning an outcome (indicated by ‘reversal’ in the graph). ‘O’ represents outcome, and ‘X’ no-outcome. (c) The structure of the probe sessions. The Probe sessions were similar to the training sessions (panel (b)), except that stage 1 actions led to the stage 2 states in a probabilistic manner. Taking action ‘L’ led to state S2 commonly (80% of the time), and to state S1 rarely (dashed lines). Taking action ‘R’ led to state S1 commonly (80% of the time), and to state S2 rarely (dashed lines).

### 2.1 Adaptive state-space representation

The stage 2 state that earned reward changed over time and, as such, subjects needed to use feedback from the previous trial to track which specific stage 2 state was rewarded so as to take the stage 1 action leading to that state.

Given this situation, it should be expected that, if a reward is earned on the previous trial, the subjects will repeat the same stage 1 action on the next trial. Figure 3a shows the odds ratio of staying on the same stage 1 action after earning a reward on the previous trial over the odds after earning no reward (across training sessions). Each bar in the graph represents a training session, and odds ratios were calculated using logistic regression analysis on the effect the reward had on staying on the same stage 1 action on the next trial (see Material and Methods for details). The horizontal dashed line in the figure shows the indifference point, i.e., when the probability of staying on the same stage 1 action after earning reward or no reward is equal. As the figure shows, in early training sessions the rats failed to show a tendency to take the same stage 1 action after earning a reward on the previous trial and instead showed a tendency to switch to the other action (first five sessions; *β* = *−*0.929 (CI: *−*1.194, *−*0.664), SE=0.135, *p <* 10^*−*11^). This sub-optimal behaviour can be explained with reference to the state-space and action representation that decisions were based on early in training; i.e., subjects were initially unaware that the environment had two stages, treated it as a single stage environment, and therefore repeated the action taken immediately prior to reward delivery. For example, if they took ‘L’ at stage 1, and ‘R’ at stage 2 and earned reward, they repeated action ‘R’ at the beginning of the next trial. Although this looks like they switched to the other stage 1 action, they were clearly repeating the last rewarded stage 2 action, which should be expected if decisions are made as if the environment is composed of a single stage.

Therefore, actions were not based on a two-stage representation, which would require that the subjects treat S1 and S2 as the outcomes of actions taken in S0, but, to the contrary, the data shows that the subjects acted as if the next action they take in S0 led to reward directly. This could be for two reasons: (i) Although S0 was visually distinct from S1 and S2, early in training it may not yet have been part of the state-space and therefore the outcomes of actions taken in S0 (which are S1/S2) were not differentially encoded from the outcomes of actions taken in S1/S2 (which were reward/no-reward). From this perspective, a reward earned by taking an action in S1 or S2 led the rats to repeat the same action in S0. (ii) S0 was part of the state-space and was being treated differently from S1/S2, but the rats had yet to encode the relationship between S0 and S1/S2. The former hypothesis relates to the state-space representation, whereas the latter is related to learning the “transition matrix” of the task, i.e., state-action-state relationships. There are two points in favour of the first hypothesis. Firstly, if animals were unsure which stage 1 action would lead to S1 (or S2), then we would expect the subjects to act randomly in S0, whereas, as the data indicate, the subject repeated the last rewarded action in S0 as if S0 was similar to the state in which the last action was taken. Secondly, animals are typically fast in learning action-outcome contingencies. For example, in a simple instrumental conditioning experiment in which two levers lead to different motivationally charged outcomes (such a food pellets), animals are able to learn the contingencies in one or two training sessions (Yin et al., 2005), whereas here it took animals more than ten training session to take the correct actions in S0. Based on this, we interpret the effect as a consequence of the simple state-space representation, i.e., all the states are organised in a single stage early in training, which is shown as S0/S1/S2 in Table 1a.

**Table 1.** Four hypotheses about state-space and action representations. (a) The state-space constitutes a single stage (outcomes such as food pellets are not shown here as states), and ‘L’ and ‘R’ are the only possible actions that the subject considers taking. (b) The state-space matches the correct state-space of the task, and the actions are ‘L’ and ‘R’. (c) The state-space only consists of state S0, and actions are ‘L → R’ and ‘R → L’. Note that single actions ‘R’ and ‘L’ are not included. (d) The state-space represents the two stages of the task and actions include both single actions and action sequences.

| representation | (a) | (b) | (c) | (d) |
| --- | --- | --- | --- | --- |
| state-space | S0/S1/S2 | S0, S1, S2 | S0 | S0, S1, S2 |
| actions | L, R | L, R | L→R, R→L | L, R, L→R, R→L |

Importantly, as Figure 3a shows, this pattern of choices reversed as the training progressed and the rats started to take the same stage 1 action that earned reward on the previous trial rather than repeating the action most proximal to reward. Replications of this finding using other experimental parameters are provided in Supplementary Experiments 1-3 in Supplementary Materials. One explanation for this observation is that, at this point in training, the rats realised that the task had two-stages and, at that point, acquired the “correct” state-space of the task (Table 1b), where the correct representation refers to the Markov Decision Process underlying the task structure (note that defining the correct state-space is not trivial; see Botvinick et al., 2015). If this is true, however, then, during the course of training, the state-space used by the animals expanded from a simple representation (Table 1a) to a more complex representation consistent with the task state-space (Table 1b).

### 2.2 Adaptive action representation

Learning the state-space of the task is not the only way that the rats could have adapted to the two-stage structure of the environment; in this task reward can be earned either by executing ‘L’ at stage 1 and ‘R’ at stage 2, or by executing ‘R’ at stage 1 and ‘L’ at stage 2. As such, it is possible that animals chunked actions ‘L’ and ‘R’ to make action sequences; say, ‘L*→*R’, and ‘R*→*L’. Using these new actions, the rats could then repeat an action sequence on the next trial after earning a reward instead of merely repeating the action proximal to the reward, as early in training. If this is true, however, then the transition in the pattern of stage 1 actions shown in Figure 2a could have been driven by the rats acquiring action sequences rather then the state-space of the task. This assumption is also consistent with the fact that the delay between the first and second action (the rats’ reaction time) decreased as training progressed (Figure 3b), a sign of the formation of action chunks.

In order to test the role of action sequences, we looked at the choices of the subjects in a *probe session* inserted at the end of the training sessions (last bar in Figure 3a), in which in a small portion of the trials transitioning between stage 1 actions and stage 2 states were switched (Figure 2c). For example, during training (non-probe sessions), after executing action ‘L’ subjects always ended up in state S2; however, the probe session included some rare transitions (20% of the trials), in which, after taking ‘L’, subjects ended up in state S1 instead of S2 (inspired by the two-stage task in Daw et al., 2011). As a consequence, in the probe session, after repeating the same stage 1 action subjects might end up in a different stage 2 state than on the previous trial, meaning that they would next take a different stage 2 action if they are selecting actions one by one. If, however, subjects are repeating the previously rewarded action sequence, we should expect them to repeat not only the first action, but also the second action even if they are now in a different stage 2 state (Keele, 1968; Dezfouli and Balleine, 2012, 2013). Figure 3c shows an example of this situation. Animals have earned reward from action sequence ‘L*→*R’ on the previous trial and have repeated action ‘L’ at stage 1 of the next trial, but on this trial have ended up in state S1 in which action ‘L’ should be taken. If, however, they take action ‘R’ this can be taken as a sign that they are repeating the whole action sequence rewarded on the previous trial.

Figure 3d shows the probability of staying on the same stage 2 action on the trials in which the stage 2 state is different from that of the previous trial. As the figure shows, if the previous trial is rewarded (‘reward’ condition) and subjects stay on the same stage 1 action (‘stay’ condition) then there is a high chance that they will also repeat the same stage 2 action, indicating that they are repeating the whole previously rewarded action sequence. This is supported by a significant interaction between staying on the same stage 1 action and reward on the previous trial (*β* = 0.494 (CI: 0.055, 0.933), SE=0.224, *p* = 0.027; see Table 2:stage 2 for the full analysis). Therefore, the pattern of choices at stage 2 is consistent with the suggestion that the subjects have expanded the initial set of actions, that previously only included actions ‘L’ and ‘R’ (Table 1a), to a more complex set that includes action sequences ‘L*→*R’ and ‘R*→*L’ (Table 1c).

**Figure 3.**
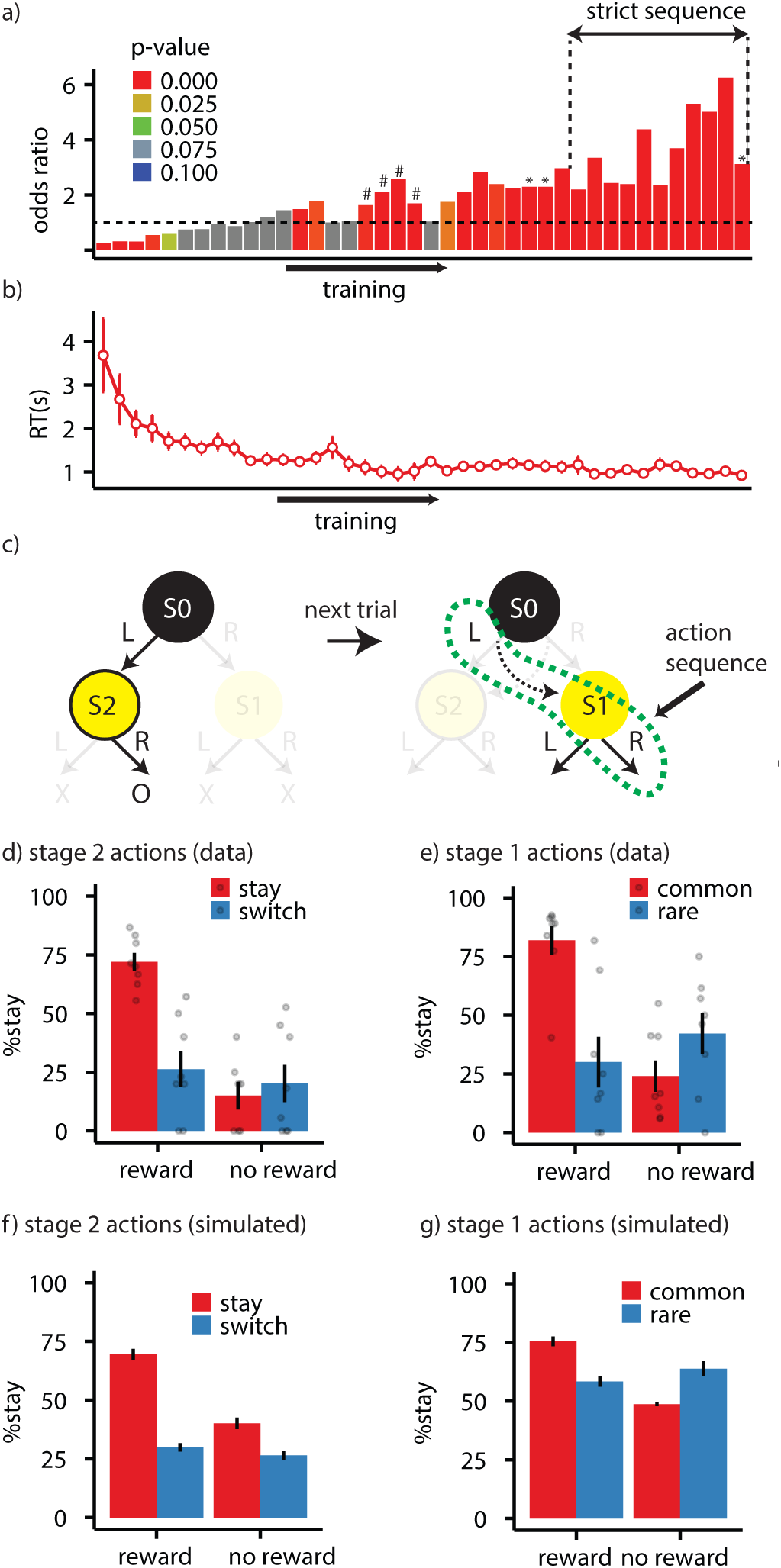
(a) Odds ratio of staying on the same stage 1 action after getting rewarded on the previous trial over the odds ratio after not getting rewarded. The dotted line represents the indifference point (equal probability of staying on the same stage 1 action after reward or no reward). Each bar represents the odds ratio for a single training session. In the sessions marked with ‘#’ in Figure 3a the contingency between stage 1 actions and stage 2 states were revered (‘L’ leads to S1 and ‘R’ to S2). ‘Strict sequence’ refers to sessions in which a trial was aborted if the animal entered the magazine between stage 1 and stage 2 actions. (b) Reaction times (RT) averaged over subjects. RT refers to the delay between performing the stage 1 and stage 2 actions. Each dot represents a training session. (c) An example of how the performance of action sequences can be detected in the probe session. On a certain trial a rat has earned a reward by taking ‘L’ at stage 1 and ‘R’ at stage 2. The subject then repeats the whole action sequence (‘L’ and then ‘R’), even though after executing ‘L’ it ends up in S1 (due to a rare transition) and action ‘R’ is never rewarded in that state. (d) The probability of staying on the same stage 2 action in the probe session averaged over subjects, as a function of whether the previous trial was rewarded (reward/no reward) and whether subjects stayed on the same stage 1 action (stay/switch). (e) The probability of staying on the same stage 1 action in the probe session averaged over subjects as a function of whether the previous trial was rewarded (reward/no reward) and whether the transition in the previous trial was common or rare. (f) Simulation of stage 2 choices, and (g) stage 1 choices using the best-fitted parameters for each subject. Error bars represent *±*1 SEM.

### 2.3 The integration of adaptive state-space and action representations

The analysis provided in the previous section showed that acquiring either the state-space representation or action sequences can explain the pattern of choices observed during the course of training (Figure 3a). Furthermore, the pattern of choices at stage 2 of the probe session provided evidence that the subjects are using action sequences. It remains open to question, therefore, whether the rats are exclusively solving the task using action sequences without relying on the state-space of the task (Table 1c), or are using both an expanded state-space and action representations for decision-making (Table 1d). To answer this question we looked at the pattern of choices at stage 1 of the probe session.

As argued in the previous sections, if decisions are based on the true state-space of the task then we expect that, after earning reward on a trial, the same stage 1 action will be taken on the next trial. The same is not true for the trials with rare transitions in the probe session, however. This is because, if the reward was earned on a trial with a rare transition, the subjects should then switch to the other stage 1 action on the next trial if they are using their knowledge of the state-space of the task (Daw et al., 2011). For example, imagine it is a trial with a rare transition and the rat, by taking ‘L’, is transferred to state S1 and earns reward. On the next trial, using the state-space of the task, the rat should switch to ‘R’ at stage 1 because ‘R’ is the stage 1 action that commonly (80% of time) leads to S1. As a consequence, staying on the same stage 1 action after earning reward depends both on the reward and the transition type on the previous trial.

On the other hand, if the rats are exclusively using action sequences without relying on the state-space of the task (Table 1c), then staying on the same stage 1 action only requires that the previous trial was rewarded; the transition type of the previous trial should not have any effect. This is because earning reward by executing an action sequence will result in the same action sequence being repeated on the next trial (and so the same stage 1 action) irrespective of the transition type on the previous trial. Therefore, a main effect of reward on staying on the same stage 1 action in the next trial indicates that the subjects are using action sequences whereas, an interaction between reward and transition type on the previous trial indicates that the subjects are using the true state-space of the task.

Importantly, the results of stage 1 actions, presented in Figure 3e, clearly revealed a significant reward-transition interaction (Table 2:stage 1), indicating that the subjects were using the correct state-space of the task (Table 1b). In addition, the main effect of reward was also significant, which indicates that the subjects were also using action sequences (Table 1c). As such the pattern of choices indicates that the rats were using both action sequences and single actions guided by the true state-space of the task. Therefore, evidence from this study suggests that, as training progressed, the initially simple state-space and action representations (Table 2a) were expanded to align with the true structure of the task (Table 2d).

**Table 2.** Results of the logistic regression analysis of stage 1 and stage 2 choices in the probe session. For the stage 1 choices, the analysis focused on staying on the same stage 1 action on the next trial, based on whether the previous trial was rewarded and whether it was common or rare (trans). ‘reward:transition’ is the interaction between reward, and transition type. For stage 2 choices, the analysis focused on staying on the same stage 2 action, based on staying on the same stage 1 action (stay) and earning a reward on the previous trial. ‘reward:stay’ is the interaction between ‘reward’, and ‘stay’.

| stage 1 actions |  |  |  |  |
| --- | --- | --- | --- | --- |
|  | intercept | reward | transition | reward:transition |
| *p*-value | 0.373 | 0.003 | 0.164 | $< 10^{-5}$ |
| $\beta$ (SE) | -0.16 (0.18) | 0.61 (0.20) | 0.37 (0.26) | 0.92 (0.20) |
| stage 2 actions |  |  |  |  |
|  | intercept | reward | stay | reward:stay |
| *p*-value | 0.001 | $< 0.001$ | 0.019 | 0.027 |
| $\beta$ (SE) | -0.95 (0.29) | 0.84 (0.23) | 0.60 (0.25) | 0.49 (0.22) |

Note that there are other explanations for the main effect of reward, other than using action sequences. For example, it could be the case that after experiencing a rare transition, subjects presumed that the relationship between stage 1 actions and stage 2 states had switched, which predicts a main effect of reward on staying on the same stage 1 action even if subjects are not using action sequences. Another explanation for the main effect of reward is based on the notion of ‘model-free’ actions. Intuitively, it implies that earning reward after taking an action increases the chance of repeating the action. In this task, it implies that reward increases the chance of taking the same stage 1 action in the next trial whether the experienced transition was common or rare (Daw et al., 2011).

Nevertheless, although these two accounts can predict a main effect of reward, they do not predict nor can they explain the effect observed on the stage 2 actions. As a consequence, we interpret the main affect of reward as indictating that the rats were using action sequences, which can explain both the stage 1 and stage 2 actions taken by the rats (see Discussion for further details).

### 2.4 Potential effects of latent states and action biases

In the previous section, we argued that the reward-transition interaction is a sign the animals had acquired an adaptive state-space representation, which allowed them to learn the relationship between stage 1 actions and state 2 states. However, recently, Akam et al. (2015) argued that this form of reward-transition interaction in multistage decision-making can be explained if subjects have learned the ‘latent states’ of the task without relying on the relationship between stage 1 actions and stage 2 states. On this account, the rats simply learned a kind of rule: e.g., whenever a reward is earned in S1, perform ‘R’ on the next trial (at stage 1), and whenever a reward is earned from S2, perform ‘L’ on the next trial. Akam et al. (2015) argue that this process requires the subjects to expand their representation of the state-space by turning S0 into latent states S0S1 and S0S2, which encode which stage 2 state was rewarded on the previous trial and, therefore, their argument depends on the rats expanding their representation of the state-space to include new states. As a consequence, even under Akam et al. (2015)’s account, the observed reward-transition interaction is evidence for an adaptive state-space representation, as we have argued. Furthermore, there are two issues for this account in the context of the current data: Firstly, it really only applies to cases where animals have been extensively trained using the rare transitions, whereas we incorporate these transitions only during probe test sessions. Secondly, this account cannot explain the adaptive action representations that we observe; i.e., the pattern of choices due to the formation of action sequences. This is because Akam et al. (2015)’s account does not imply repeating the previously rewarded sequence of actions in S0 and S1/S2.

There is another potential interpretation of the reward-transition interaction based on the potential for a local response bias induced by the reward function. Assume that, in a part of the probe session, actions taken in S1 are rewarded (and actions taken in S2 are not), and by trial and error the animal develops a tendency to take action ‘R’ more frequently than ‘L’ at S0; i.e., the probability of staying on the same action when it is ‘R’ is higher than when it is ‘L’. As most of the common transitions after taking ‘R’ are rewarded (as they mostly lead to S1) and most of the rare transitions are non-rewarded (as they mostly lead to S2), there will be an effect of reward-transition interaction on the probability of staying on the same action, which looks like the animals are taking the structure of the world into account, while what they are doing is simply taking action ‘R’ more frequently. This issue was discussed in Dezfouli and Balleine (2013); Smittenaar et al. (2013) and further analysed in Akam et al. (2015) and one way to address it is to add a new predictor to the analysis of the effect of reward and transition on staying on the same stage 1 action. This new predictor encodes whether the previous stage 1 action was the best action, i.e., it leads to the stage 2 state with the highest reward, which will absorb the effect of the reward-transition interaction if the interaction is just due to repeating the best action more frequently (Smittenaar et al., 2013; Akam et al., 2015). This analysis is presented in Table A4, which shows that even in the presence of this predictor the effect of reward and the reward-transition interaction are still significant. As such, the reward-transition interaction is unlikely to be due to this form of response bias.

### 2.5 Choice of experimental parameters

Animals were given three probe sessions in total, and the results reported above were taken from the last of these tests which was the final experimental session (probe sessions are marked by an asterisk in Figure 3e). The full analysis of all the probe sessions is presented in see Table A3 in Supplementary Materials. The structure of the probe sessions was identical to each other and also in terms of results; similar to the third probe session analysed above, the main effect of reward was significant in sessions one and two. However, unlike the last probe session, in the first two sessions, the rats did not show evidence that they were using action sequences (Table A3: probe 1 and 2, stage 2 actions; reward-stay interaction; *p*-value>0.1). A closer examination of these sessions revealed that, at this stage in the training, the rats were not discriminating between the stage 2 stimuli; i.e., because the analysis of stage 2 only included trials in which the stage 2 state is different from the previous trial, we expected the probability of staying on the same stage 2 action to be generally low (as different actions are rewarded in the stage 2 states), which was not the case in the first two probe sessions (see Table A3; *p*-value *>* 0.05 for the intercept term at stage 2 actions in probe 1, 2). As a consequence, under these conditions, staying on the wrong stage 2 action due to the performance of action sequences cannot be detected, because the rats are likely to take the incorrect action at stage 2 states even if they are not taking an action sequence. One reason for this lack of discrimination is the potential for interference between the stage 1 and stage 2 actions; if the rats checked the magazine after taking the stage 1 action they may then have repeated the same stage 1 action instead of taking the correct stage 2 action. This would make it look like the animals were not discriminating between stage 2 stimuli. To address this issue we introduced the ‘strict sequences’ criterion for the next ten training sessions (Figure 3e) under which a trial was aborted if a rat entered the magazine between the stage 1 and stage 2 actions. After these ten training sessions the rats were given the third probe test, in which they showed they were able significantly to discriminate between the stage 2 states (see Table 2; *p*-value = 0.001 for the intercept term at stage 2 actions). Note that the analysis presented in the previous and subsequent sections relates to this last probe session.

In Supplementary Materials, we also present three supplemental experiments each of which used different parameters. Supplementary experiment 1, which is shown in Figure A3, includes probe sessions throughout the training process and shows the development of choices. Supplementary experiments 2 and 3 are mostly similar to each other and provide a training protocol in which animals reliably exhibit the reward effect in their stage 1 actions. However, in none of these experiments were we able to observed the performance of action sequences, as indicated by the reward-same interaction in stage 2 actions (see Table A5 for the full analysis of supplementary experiment 1 and Table A6 for the full analysis of supplementary experiments 2,3). One main difference between these experiments and the experiment reported in the main paper is that, whereas in the main experiment the ITI was zero, in the supplementary experiments, the inter-trial interval was non-zero. In this latter condition, animals were often found to take actions during the ITI, which were not rewarded but which were very likely to interfere with the performance of action sequences once the next trial started. Using an ITI of zero addressed this issue. Lastly, as Figure 3a shows, there were some training sessions in which the contingency between stage 1 actions and stage 2 states was reversed. These training sessions were introduced to overcome the interference that has been argued to be produced when animals, including humans, are first exposed to rare trails.

### 2.6 Computational models of adaptive decision-making

We next sought to establish the computational model that best characterized the decision-making process used by the rats in this experiment. The modelling was focused on the probe session that we analysed in the previous sections (the final probe session). For this purpose, we compared different families of reinforcement-learning (RL) model to establish which provided a better explanation for the data (Table 3). The families compared included: (1) a non-hierarchical model-based RL family (MB) corresponding to Table 1b, which assumes that the subjects acquired the correct state-space of the task, but in which action sequences were *not* included in the set of actions; (2) a hierarchical RL family (H) corresponding to Table 1c, which assumes that the set of actions included only action sequences, but that decisions were *not* guided by the true state-space of the task; (3) a hierarchical model-based RL family (H-MB) corresponding to Table 1d, which assumes that the subjects were using single actions, action sequences and the true state-space representation for decision-making (Dezfouli and Balleine, 2013); (4) a model-free RL family (MF), and (5) a hybrid model-based RL and model-free RL family (MB-MF). These latter two families have been previously used to characterise performance on a similar task (Daw et al., 2011), and we used them as baselines.

In total we considered 344 different models in which each family consisted of several members with different degrees of freedom (see Supplementary Materials for details). We then calculated the negative log model-evidence for each model *M* given the choices of subjects, *D* (denoted by *−* log *p*(*D|M*)). Table 3 shows the negative log model-evidence along with other properties of the best model for each family. The differences in log model-evidence (log-Bayes factor) between the best fitting model of the H-MB family and other families was greater than 13. In the Bayesian model comparison literature, log-Bayes factors greater than 3 are considered to be strong evidence (Jeffreys, 1961). Therefore, the above results provide strong evidence that the subjects were utilising H-MB to guide action selection. Figure 4 shows the negative log model-evidence for the best eight models in each family and shows that the different members of the H-MB family provide a better explanation of the data than any of the other families.

**Table 3.** States and actions for each family of computational model along with degrees-of-freedom (df), pseudo-*r*^2^, and negative log model-evidence ( log *p*(*D M*)) for the best model in each family; lower values of negative log model-evidence are better.

| family | state-space | actions | df | p-$r^2$ | negative log model-evidence |
| --- | --- | --- | --- | --- | --- |
| MB | S0, S1, S2 | L, R | 5 | 0.175 | 1222.354 |
| H | S0 | L→R, R→L | 4 | 0.207 | 1185.274 |
| H-MB | S0, S1, S2 | L, R, R→L, L→R | 4 | 0.207 | 1172.087 |
| MF | S0, S1, S2 | L, R | 4 | 0.174 | 1219.575 |
| MB-MF | S0, S1, S2 | L, R | 5 | 0.179 | 1223.023 |

**Figure 4.**
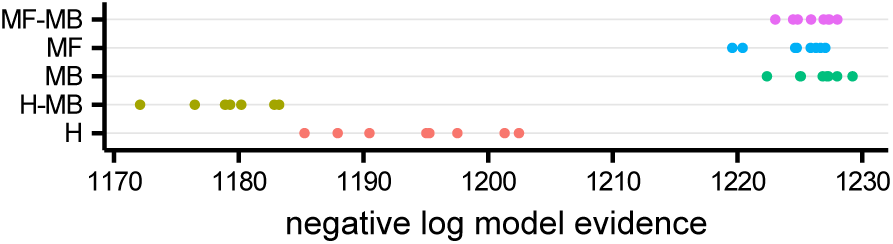
Negative log model-evidence ( log *p*(*D M*); lower numbers indicate better models) for the first best eight models in each family of computational models.

We then simulated eight instances of the H-MB model of the task using the best fitting parameters for each subject (Table A2) and analysed the stage 1 and stage 2 choices of the simulated model. Analysis of stage 1 choices (Figure 3f) revealed a significant main effect of reward (*β* = 0.255 (CI: 0.191, 0.320), SE=0.033, *p <* 10^*−*15^), and a significant interaction between whether the previous trial was rewarded and the transition type of the previous trial (*β* = 0.291 (CI: 0.193, 0.388), SE=0.049, *p <* 10^*−*8^). Analysis of stage 2 choices (Figure 3g), revealed a significant interaction between earning a reward on the previous trial and the likelihood of staying on the same stage 1 action (*β* = 0.259 (CI: 0.166, 0.353), SE=0.047, *p <* 10^*−*7^). These results are, therefore, entirely consistent with the behavioural results of our experiments using rats as subjects. Furthermore, the fact that H-MB family provides a better fit than the H family implies that subjects were using the correct state-space of the task (MB part), and the H-MB family being better than MB family implies that subjects were using action sequences. The H-MB family also provided a better fit than baseline MB/MF models, however, this does not imply that some form of model-free RL is not working concurrently with a H-MB model, as we discuss in the next sections.

## 3 Discussion

Learning the value of different actions in various states of the environment is essential for decision-making in multi-stage environments. This learning process operates above the state-space and action representation and, therefore, the ability to (i) acquire the correct state-space of the task, and (ii) create new actions that are useful for solving the task, are important for efficient decision-making. Using a sequential decision-making task in rats, we provide direct evidence that, early in training, subjects make decisions based on simple state-space and action representations, but that, during the course of training, both state-space and action representations evolve and adapt to the structure of the environment. We found that this adaptive process included (i) expanding the state-space representation to include multiple stages of the environment, and (ii) augmenting the set of actions with new action sequences that are useful for earning rewards.

The ability to solve multi-stage decision-making tasks has been previously demonstrated in different species, however, unlike the current study, these demonstrations have either given the different stages of the task to the subjects (e.g., Daw et al., 2011), or have explicitly signalled the actions that should be taken at each stage (e.g., Christie and Dalrymple-Alford, 2004), which remove the necessity for building multi-stage representations to solve the task. Similarly, the ability of animals to concatenate simple actions to execute action sequences has been established previously and here we extended these prior studies by showing that, during the course of learning, single actions turn into action sequences that are not only executed, but also are evaluated as a single response unit (Ostlund et al., 2009).

A task similar to the the two-stage task that we used here in rats has previously been employed to study different forms of decision-making processes in humans (Daw et al., 2011). Although the experiments in those studies were composed of a single session, results indicated that the subjects were using an expanded state-space representation without needing to go through multiple training sessions. This is presumably due to the instructions and the cover story provided to the subjects, which informed them about the correct state-space representation. In terms of acquiring action sequences, using a similar task in humans we have previously shown that subjects engaged action sequences (Dezfouli and Balleine, 2013). Again, however, we found they were able to do so without requiring multiple training sessions. Why such sequences should have formed so rapidly is a matter of conjecture but, as the task involved pressing keys on a keyboard, familiarity with similar response sequences could have supported sequence acquisition (especially as only two key presses were required to earn the reward). Based on these comparisons, the results of the current experiments point to the importance and complexity of learning state-space and action representations. As the profile of the the rats’ choices indicates, they required a significant amount of training in order to learn the structure of the environment (10-40 sessions). This is while, in some instrumental conditioning settings, animals are able to learn the contingency between actions and states in two and sometimes in a single training session (Yin et al., 2005). Uncovering the processes that determine the encoding of the state-space of the task and how this process interacts with that subserving instrumental conditioning will be an important step towards better understanding the learning mechanisms that mediate decision-making processes generally.

The results of the computational modelling indicated that hierarchical model-based RL provides the best explanation for the rats’ choices. This model assumes that the subjects build an internal map of the environment which encodes both the specific outcomes of single actions *and* of action sequences. The validity of this assumption for single actions can be confirmed based on the results of the current experiment and previous studies (Balleine, 2005) showing that subjects encode the specific outcome of each individual action, e.g., taking ’R’ leads to ’S1’ and ‘L’ leads to ’S2’. With regard to encoding the outcome of action sequences, although previous studies have indicated that the subjects specifically encode the outcome of each action sequence (Ostlund et al., 2009), we cannot assess whether subjects encoded outcome specific sequences in the current study because the value of the food outcome was not manipulated. As such, the results do not address the (model-based or model-free) nature of the controller mediating the evaluation of action sequences.

In the same line, as we discussed in the previous sections, the emergence and detection of action sequences requires certain experimental conditions, such as a short ITI. Without these conditions, actions at stage 1 are consistent with the operation of action sequences, but not actions at stage 2. One explanation for this effect could be the potential inhibition of action sequences. For example, since during long ITIs subjects go through extinction, it is unlikely that they keep performing the whole action sequence throughout the ITI and in the next trial. As such, although the first component of the action sequences at stage 1 is performed, the second component is inhibited when it is inappropriate, making it harder to detect the performance of action sequences. Alternatively, this pattern of choices could also indicate the operation of another RL system, such as model-free RL, instead of interrupted sequences. Within this alternative framework, choices are a mixture of goal-directed actions (model-based), and model-free actions that are guided by their ‘cached’ (as opposed to their current values; Daw et al., 2005). Our results are ambivalent with respect to this interpretation, but since, in other conditions, there is positive evidence for action sequences, it is more parsimonious to interpret this result in terms of the inhibition of action sequences rather than being the output of an additional model-free system.

## 4 Methods and Materials

### 4.1 Subjects and apparatus

Eight experimentally naive male Hooded Wistar rats served as subjects in this study. Data from all the subjects are included in the analyses. All animals were housed in groups of two or three and handled daily for one week before training. Training and testing took place in eight Med Associates operant chambers housed within sound- and light-resistant shells. The chambers were also equipped with a pellet dispenser that delivered one 45 mg pellet when activated (Bio-Serve). The chambers contained two retractable levers that could be inserted to the left and the right of the magazine. The chambers contained a white noise generator, a Sonalert that delivered a 3 kHz tone, and a solenoid that, when activated, delivered a 5 Hz clicker stimulus. All stimuli were adjusted to 80 dB in the presence of a background noise of 60 dB provided by a ventilation fan. A 3 W, 24 V house light mounted on the wall opposite the levers and magazine illuminated the chamber. Microcomputers equipped with MED-PC software (Med Associates) controlled the equipment and recorded responses. Animals were food deprived one week before starting behavioral procedures. They were fed sufficiently to maintain their weight at 90% of their free-feeding weight. The animals were fed after the training sessions each day and had free access to tap water whilst in their home cage. Each training session (except the magazine training sessions) started with insertion of the levers, and ended with their retraction. All procedures were approved by the University of Sydney Animal Ethics Committee.

### 4.2 Behavioral procedures

Rats were given two sessions of magazine training in which 30 grain pellets were delivered on a random time 60-s schedule (Figure 1:phase 1). Rats were then trained to lever press on a continuous reinforcement schedule with one session on the left lever and one session on the right lever each day for four days with the total number of outcomes each day limited to 60 per session (Figure 1:phase 2). The total duration of each session was limited to 60 minutes. Next, rats were trained to discriminate the two stimuli (Figure 1:phase 3). Each session started with the presentation of a stimulus. The stimulus was presented until the rat performed an action (either pressing the left or right lever) after which the stimulus turned off. For one stimulus, taking the left action led to the reward, whereas for the other stimulus taking the right action led to reward. Levers and stimuli were counterbalanced across subjects. After an action was chosen, there was a 60-second inter-trial interval (ITI) after which the next trial started with the presentation of the next stimulus, again chosen randomly. The duration of each session was 90 minutes, with no limit on the maximum number of earned rewards. The stimuli were a constant or a blinking house light (5 Hz). The result of this phase is depicted in Figure A2.

The rats then received training on the two-stage task depicted in Figure 2b (maximum 60 outcomes in a session and maximum duration of a session was limited to 1 hour). Animals were trained on the two-stage task for 40 sessions. In the middle of, or at the end of these training sessions, they were given probe sessions, similar to the training sessions except that stage 1 actions led to stage 2 states in a probabilistic manner (Figure 2c). These sessions are indicated by ‘*’ in Figure 3a. After the first two training sessions, subjects then received ten more training sessions, and were then given a further probe test. The results reported in the Results section correspond to this last probe session. In the sessions marked with ‘#’ in Figure 3a the contingency between stage 1 actions and stage 2 states were revered (‘L’ leads to S1 and ‘R’ to S2); these training sessions were followed by a session in which ‘L’ leads to S1 and ‘R’ to S2 in 20% of times, followed by normal training sessions, as Figure 3a shows. Finally, ‘strict sequence’ in Figure 3a refers to a session in which a trial was aborted if the animal entered the magazine between stage 1 and stage 2 actions. In all the training phases levers were present throughout the training session.

### 4.3 Behavioral analysis

We used R (R Core Team, 2016) and lme4 packages (Bates et al., 2015) to perform a generalized linear mixed effects analysis. In all of the analyses, logistic regression was used and all the fixed effects (including intercepts) were treated as random effects varying across subjects. For analyses that included more than one session, random effects were assumed to vary across sessions and subjects in a nested manner. Confidence intervals (CI) of the estimates were calculated using the ‘confint’ method of lme4 package with the ‘Wald’ parameter.

In the analyses of the stage 1 of non-probe sessions, we used a logistic regression analysis in which the independent predictor was whether the previous trial was rewarded (reward or no-reward), and the dependent variable was staying on the same stage 1 action. The *p*-value of this analysis was used in Figure 3a for colour-coding each bar, and the height of each bar represented the odds ratio calculated as *e*^*β*^. The intercept term of this analysis is shown in Figure A1. In the analyses of stage 1 of the probe sessions, the independent predictors were transition type of the previous trial (rare or common) and whether the previous trial was rewarded (reward or no-reward), whereas the dependent variable was staying on the same stage 1 action. The effects of interest were reward and the reward by transition-type interaction. In the analysis of stage 2 probe sessions, the independent variables were whether the stage 1 action was repeated (same stage 1), and whether the previous trial was rewarded. The dependent variable was staying on the same stage 2 action. The effect of interest was the interaction between the two independent variables. Note that only trials in which the stage 2 state was different from the stage 2 state of the previous trial were included in this analysis.

In all the analyses, only trials in which subjects made a correct discrimination on the previous trial (‘R’ in S2, and ‘L’ in S1) were included (%71 of trials in the whole training period). This was for two reasons. Firstly, it was not clear how subjects learn from actions taken during incorrect discriminations, which were never rewarded. Secondly, as depicted in Figure 3c, for the analysis of adaptive action representation, we focused on the trials in which the stage 2 states was different from that of the previous trial. When executing action sequences, we expected the subject to take the same stage 2 action in the next trial if (i) they were rewarded in the previous trial and (ii) they take the same stage 1 action, but not otherwise (as we focused on consecutive trials with different stage 2 states). However, assume that the subject makes an incorrect discrimination in the previous trial, e.g., it takes action ‘L’ at stage 1, moves to state S2 and takes action ‘L’ in that state, which is not rewarded since action ‘L’ in S2 is never rewarded. In the next trial, if the subject takes action ‘L’ again and ends up in state S1 (Figure 3c), there is a high chance that it will take ‘L’ again at stage 2, since ‘R’ is never rewarded in S1. Therefore, even if no reward was earned in the previous trial, there is a high chance that the subject will repeat the same stage 2 action in a different trial. This only happens in the condition that the subject made an incorrect discrimination in the previous trial, and in order to remove this interaction between the discrimination between actions at stage 2, and the analysis of action sequences, we only included the trials in which the subjects made correct discrimination in the previous trial. The analysis similar to the one presented in Table A3 without removing these trials is presented in Table A4, which shows that the main statistical tests that we used to argue for adaptive state-space and action representations are statistically significant whether we include all of the trials or not.

#### 4.4 Computational modelling

Reinforcement-learning models considered for behavioural analysis were similar to the hierarchical RL models provided in Dezfouli and Balleine (2013) and the model-based/model-free family provided in Daw et al. (2011). In addition to these families, we also considered a family of hierarchical models (corresponding to the H family) in which only action sequences were available at stage 1 (i.e., single actions ‘L’ and ‘R’ were not available). In addition to the free-parameters mentioned in previous work, we added two new parameters here. The first free-parameter only applies to the hierarchical families (H, H-MB), which represented the probability that the performance of action sequences is interrupted in the middle of the action sequence (i.e., subjects only perform the first component of an action sequence, and select a new action at stage 2). The other free-parameter coded the tendency of animals to take the discriminative action at stage 2, irrespective of the value of each action (tendency to take ‘R’ in S2 and ‘L’ in S1). This free-parameter allowed the model to learn that one of the actions in each of the stage 2 states was never rewarded. Details of the computational models along with their mathematical descriptions are presented in Supplementary Materials. For the purpose of model comparison, we generated different instances of each family of models with different degrees of freedom (see Supplementary Materials for details). Model-evidence, reported in Table 3 and Figure 4 was calculated similar to Piray et al. (2014).

## 5 Acknowledgements

AD and this research were supported by grants DP150104878, FL0992409 from the Australian Research Council to BWB. BWB was supported by a Senior Principal Research Fellowship from the National Health & Medical Research Council of Australia, GNT1079561.

## Supplementary Materials

### A1 Computational modelling

We compared five different families of the RL algorithms in order to evaluate which one provides a better explanation for data. These five families are: (1) a non-hierarchical model-based RL family (MB); (2) a hierarchical RL family (H); (3) a hierarchical model-based RL family (H-MB); (4) a model-free RL family (MF), and (5) a hybrid model-based RL and model-free RL family (MB-MF). Each family had several instances, that we described them below.

We assumed that the environment has five states; the initial state denoted by S_0_, stage 2 states denoted by S_1_ and S_2_, the reward state denoted by S_Re_ and no-reward state denoted by S_NR_. For the case of non-hierarchical models, we assumed that actions L and R are available in states S_0_, S_1_ and S_2_, and for the case of hierarchical models, we assumed actions L, R, LR, LL, RL, and RR are available in states S_0_, and actions L and R, are available in states S_1_ and S_2_.

#### A1.1 Model-based RL (MB)

Model-based RL (MB) is suggested to be the computational substrate for goal-directed decision-making. The model-based system works by learning the model of the environment, and then calculating the value of actions using the learned model. The model of the environment is composed of the transition function (*T* (.)), and the reward function (*R*(.)). We denote the transition function with *T* (*s’|a, s*) which is the probability of reaching state *s*^*I*^ after executing action *a* in state *s*. We assume that the transition function at the first stage is fixed,

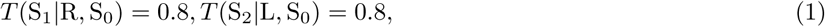

and it will not change during learning. For the other states, after executing action *a* in state *s* and reaching state *s*^*I*^, the transition function updates as follows:

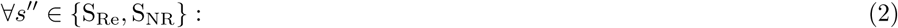

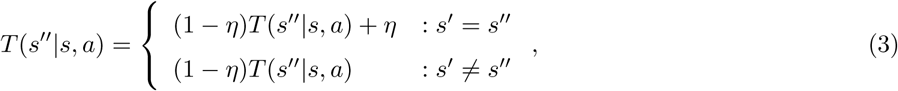

where *η*(0 *< η <* 1) is the update rate of the state-action-state transitions. For the reward functions, we assumed that the reward at state S_Re_ is one, and zero in all other states,

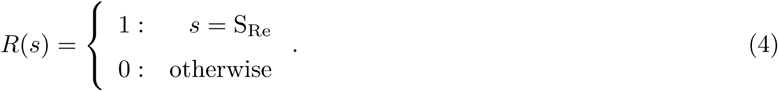

Based on the above reward and transition functions, the goal-directed (model-based) value of taking action *a* in state *s* is as follows:

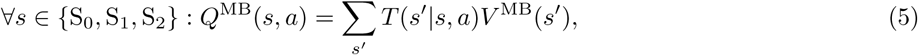

where:

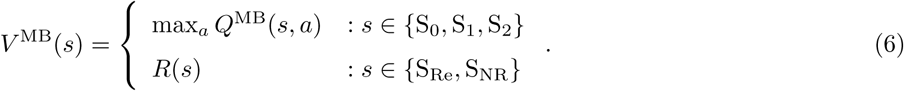

Finally, the agent uses the calculated values to choose actions. The probability of selecting action *a* in state *s*, denoted by *π*(*s, a*), will be determined according to the soft-max rule:

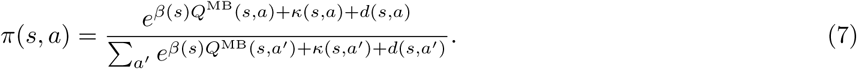

The above equation reflects the fact that actions with higher values are more likely to be selected. The *β*(*s*) parameter controls the rate of exploration; the parameter *κ*(*s, a*) is the action preservation parameter and captures the general tendency of taking the same action as the previous trial Ito and Doya (2009); Lau and Glimcher (2005).

Finally, the term *d*(*s, a*) represents the tendency of the subjects to take the discriminative actions at the stage 2 states (taking action R in S_2_ and action L in S_1_). A positive value for this parameter entails that a subject has a tendency to take the discriminative action at stage 2 states. Please note that the effect of this parameter is on top of the effect of values of the actions at the stage 2 states.

For the exploration parameter, we assume that *β*(*s*) = *β*_1_ if *s* = S_0_ and *β*(*s*) = *β*_2_ if *s ∈ {*S_1_, S_2_*}*. For the perseveration parameter we assumed that if *s* = S_0_ and *a* being the action taken in the previous trial in the S_0_ state, then *κ*(*s, a*) = *k*, otherwise it will be zero. Finally, for the discrimination parameter, we assume

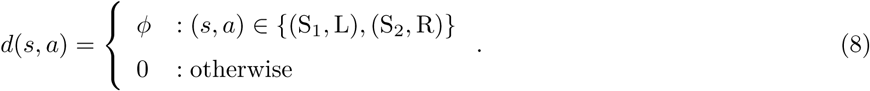

In the most general form, all the parameters (*β*_1_*, β*_2_*, η, k, φ*) were treated as free parameters. We also generated eight variants by (1) setting *β*_1_ = *β*_2_ (i.e., rate of exploration at stage 1 and stage 2 states are the same), (2) setting *k* = 0 (there is no tendency to perseverate on the previously taken actions), and (3) setting *φ* = 0 (there is no tendency to take the discriminative action at stage 2).

#### A1.2 Hierarchical model-based RL (H-MB)

Implementation of the hierarchical structure is similar to hierarchical RL, with action sequences (LL, LR, etc) as *options* (equivalent to action sequences in this setting). We assumed actions L, R, LL, LR, RL, and RR are available in stats S_0_, and actions L and R, are available in states S_1_ and S_2_. After reaching a terminal state (S_Re_ or S_NR_), transition functions of both the action sequence, and the single action that led to that state update according to equation 3. In the case of single actions, the transition function will be updated by the *η* = *η*_1_ update rate, and in the case of action sequences, the transition function will be updated by the *η* = *η*_2_ update rate. Based on the learned transition function, value of actions in each state is calculated by the goal-directed system using equation 5. Using the state-action values (*V* ^MB^(*s, a*)), the probability of selecting each action under goal-directed control will be as follows:

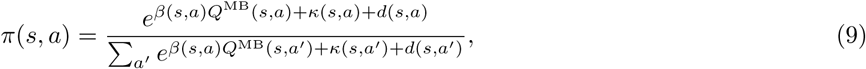

where *β*(*s, a*) is the rate of exploration. The rate of exploration for stage 2 actions (*s ∈* {*S*_1_, *S*_2_}) is *β*(*s, a*) = *β*_2_. For stage 1 actions (*s* = *S*_0_), if *a* is a single action, we assume *β*(*s, a*) = *β*_1_, and if *a* is an action sequence *β*(*s, a*) = *β*_3_:

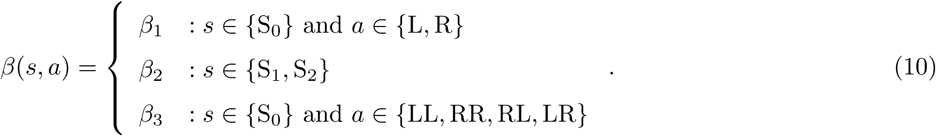

As before, *κ*(*s, a*) captures action perseveration. We assumed that *κ*(*s, a*) = *k*_1_ if action *a* is a single action, and *κ*(*s, a*) = *k*_2_ if action *a* is an action sequence:

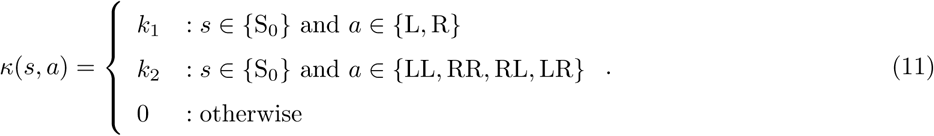

Parameter *d*(*s, a*) is similar to the one defined in the previous section.

For calculating the probability of selecting each action, equation 5 was used in the case of stage 1 actions, as these actions are chosen using using model-based valuation. In the case of stage 2 actions, however, the probability of selecting actions depends on whether the action to be executed is part an action sequence selected earlier, or is it a single action selected at stage 2 based on the model-based valuations at this stage, i.e., if the action selected at stage 1 is a single action then the action at stage 2 will be selected using equation 5, however, if the action selected at stage 1 is an action sequence, then the second component of the selected action sequence will be executed in stage 2. Based on this, we calculated the probabilities of selecting actions at stage 2 as follows. Assume we know action L has been executed in state S_0_ by the subject; then, the probability of this action being due to performing the LR action sequence is:

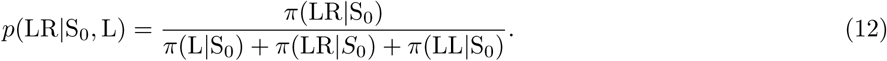

Similarly, the probability of observing L due to selecting the single action L at stage 1 is:

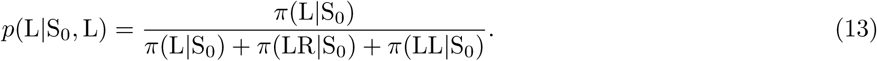

Based on this, the probability that the model assigns to action *a* in state *s ∈* {S_1_, S_2_}, given that action *a*^*I*^ is being observed in S_0_ is:

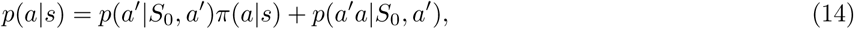

where 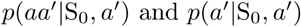 are calculated using equations 12 and 13 respectively.

Next, we assumed that even under the conditions in which an action sequence is being executed, there is a chance that the performance of the action sequence will be interrupted at stage 2, that is, a subject selects an action sequence at stage 1, but stops executing the action sequence at stage 2, and selects a new action using model-based evaluations (equation 5). This variant is inspired by similar approaches in the hierarchical RL literature (see Hengst (2012) for a review).

Let’s assume that the probability of interrupting an action sequence is *I*, then equation 14 will become as follows:

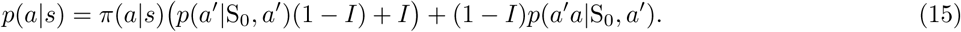

It can be verified that in the case of *I* = 0, i.e., action sequences never become interrupted, the above equation will degenerate to equation 14. In the case of *I* = 1, i.e., all the action sequences are interrupted and they have no effect on stage 2 choices, and we have:

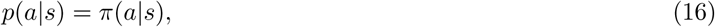

which indicates that the probability of taking each action at stage 2 is guided only by the rewards earned on that stage 2, and not by the action sequences in the first stage.

In the most general form, all the free parameters are included in the model: *β*_1_, *β*_2_, *β*_3_, *η*_1_, *η*_2_, *k*_1_, *k*_2_, *φ*, *I*. We generated 256 simpler models by setting (1) *β*_1_ = *β*_2_ (exploration rates at stage 1 and stage 2 choices are the same), (2) *β*_1_ = *β*_3_ (exploration rates for action sequences and single actions are the same), (3) *η*_1_ = *η*_3_ (learning rates for action sequences and single actions are the same), (4) *k*_1_ = 0 (no perseveration for single actions), (5) *k*_2_ = 0 (no perseveration for action sequences), (6) *φ* = 0 (no tendency to take discriminative actions), (7) *I* = 1 (action sequences are always interrupted), (8) *I* = 0 (action sequences are never interrupted).

#### A1.3 Hierarchical (H)

This family is similar to H-MB family, except that only action sequences (LL, LR, RL, RR) can be selected at stage 1 (S_0_). In the most general form the free-parameters included *β*_1_ (exploration parameter at stage 1), *β*_2_ (exploration parameter at stage 2), *η*_1_ (learning rate for action/action sequences), *k*_2_ (preservation on action sequences), *φ* (discrimination parameter), *I* (sequence interruption parameter). We then generated 16 different variants by setting *β*_1_ = *β*_2_, *k*_2_ = 0, *φ* = 0, *I* = 0.

#### A1.4 Model-free RL (MF)

We used *Q*-learning (Watkins, 1989) for model-free learning. After taking action *a* in state *s*, and reaching state *s*^*I*^, the model-free values update as follows:

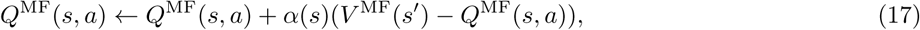

where *α*(*s*)(0 *< α*(*s*) *<* 1) is the learning rate, which can be different in stage 1 and stage 2 states,

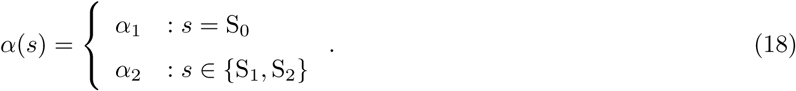

In equation 17, *V* ^MF^(*s*) is the value of the best action in state *s*:

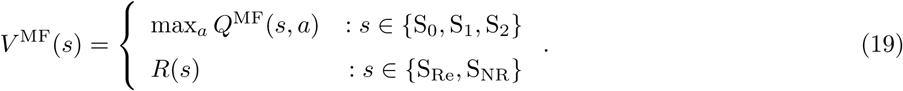

In addition to the update in equation 17 after taking actions at stage 2, the value of the action that was taken at stage 1 will also get updated according to the outcome. Assume *a* is the action that was taken in S_0_, *a*^*I*^ is the action that was subsequently taken in *s* (second stage states, i.e., *s ∈ {*S_0_, S_1_*}*), and *s*^*I*^ is the state that was visited after executing 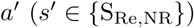 Then, action values update as follows:

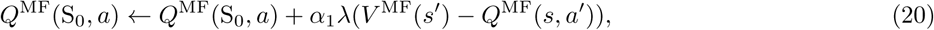

where *λ*(0 *< λ <* 1) is the reinforcement eligibility parameter, and it determines the extent to which the first stage action values are affected by receiving the outcome after executing the second stage actions. The action selection method, and variants of this form of learning are described in the next section.

#### A1.5 Model-free, model-based hybrid RL (MF-MB)

This model is a combination of model-free RL, and model-based RL, in which final action values are computed by combining the values provided by model-free and model-based processes,

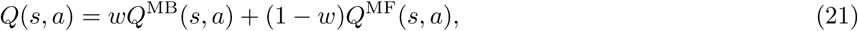

were *w*(0 *< w <* 1) determines the relative contribution of model-free and model-based values into the final values.

The probability of selecting action *a* in state *s* will be determined according to the soft-max rule:

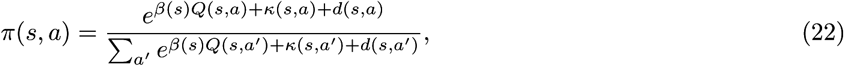

where parameters are same as the ones we described in the Model-based RL (MB) section.

In the most general form, all the free parameters are included in the model: *β*_1_, *β*_2_, *η*, *α*_1_, *λ*, *w*, *k*, *φ* (we assumed that *α*_2_ = *η*). We generated 32 simpler models by setting (1) *λ* = 0, (2) *α*_1_ = *α*_2_ (learning rate of model-free system is the same at stage 1 and stage 2 states), (3) *β*_1_ = *β*_2_ (rate of exploration is the same at stage 1 and stage 2 states), (4) *k* = 0 (there is no tendency to perseverate on the previously taken action), and (5) *φ* = 0 (there is no tendency to take the discriminative action at stage 2).

By setting *w* = 0 the above hybrid model degenerated to a model-free process described in the previous section, and therefore, we generated 32 variants of model-free RL (similar to the hybrid model), by setting *w* = 0.

#### A1.6 Model comparison

We took a hierarchical Bayesian approach to compare different models. This approach provides a framework to compare models based on their complexity and their fit to data. Bayesian model comparison is based on the model evidence quantity, which is the probability of the data given a model. The approach that we took to calculate this quantity is similar to the approach taken in Piray et al. (2014).

For each model, there are two sets of free parameters: group-level parameters denoted by Θ (we call these parameters hyper-parameters), and subject-level parameters, denoted with *θ*_*i*_ for subject *i*. The hyper-parameters define the prior distribution over subject-level parameters. The aim is to calculate the probability of data (denoted by *D*) given model *M*:

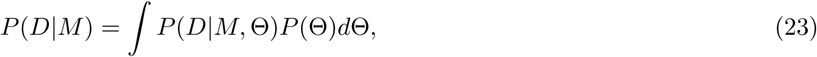

Since the above integral is intractable, we approximate it using Bayesian Information Criterion (BIC) (Schwarz, 1978):

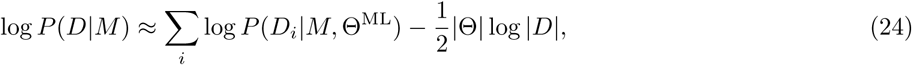

where Θ^ML^ is the maximum-likelihood estimate of Θ, and *D*_*i*_ is the data of subject *i*. *|*Θ*|* is the number of hyper-parameters, and *|D|* is the sum of number of choices made by all the subjects. In the above formula, the term inside the sum is:

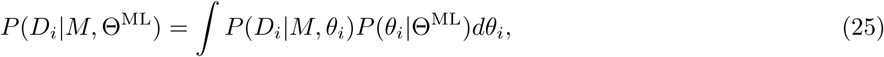

which is again intractable to compute, and we use Laplace method (MacKay, 2003) to approximate it:

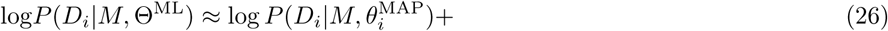

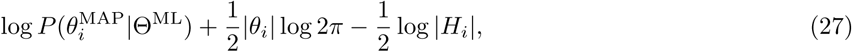

where 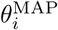 is the maximum a posterior (MAP) estimate of *θ*_*i*_. |*θ*_*i*_| is the number of free parameters for model *M*, and |*H*_*i*_| is determinant of the Hessian matrix at 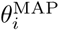.

Thus in summary, we calculated the model evidence for each subject using equation 27, and then we summed over all the model evidence for all the subjects to calculate equation 24, which is the model evidence over the whole group.

The only remaining question is how to calculate Θ^ML^, which is:

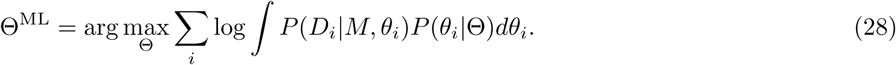

Similar to Huys et al. (2011), we solved the above optimization problem using the expectation-maximization (EM) procedure (Dempster et al., 1977). This procedure starts with an initial value for the hyper-parameters Θ, using which the posterior distribution of each individual’s parameter will be estimated using the Laplace approximation. These individual posterior distributions then shape a new value for the hyper-parameters (Θ), which will be used again to get new posterior distribution for each individual. This process continues until the hyper-parameters do not change anymore across iterations. Please refer to Huys et al. (2011) for the details of the method.

The prior over all the individual level parameters (*θ*_*i*_) where assumed to be a Gaussian distribution, and the mean and variance of the Gaussian were included in the hyper-parameters (thus the number of hyper-parameters for each model were twice as the number of free parameters of the model). Parameters that had a limited range (e.g., learning rates), were transformed to stratify the constrains.

We used the NLOPT software package (Johnson,) for nonlinear optimization using ‘BOBYQA’ algorithm. Finally, we used ‘DerApproximator’ package (Kroshko,) in order to estimate the Hessian at the MAP point.

#### A1.7 Model comparison results

In total we tested 344 models (H-MB:n=256, H:n=16, MB:n=8, MF:n=32, MB-MF:n=32). Table A1 shows the different properties of the best four models in each family. Out of 344 models that we tested, one of the models, which had eight free parameters (*β*_1_*, β*_3_*, η*_1_*, η*_2_*, k*_1_*, k*_2_*, φ, I*), was not identifiable (the estimated Hessian matrix was not a positive-definite matrix), and therefore it was excluded from the analysis. Table A2 represents estimated parameters for each individual in the best model (indicated by * in Table A1). The term ‘*−* log *P* (*D*_*i*_|*M,* Θ^*ML*^)’ represents the negative log-model evidence for each subject, obtained from equation 27.

#### A1.8 Supplementary experiments

The aim of this section is to present the results of three supplementary experiments (1-3) conducted using different parameters to the ones used for the experiment reported in the main paper. Each experiment shows that similar to Figure 3a there was a shift in the pattern of choices from simple representations to more complex representations that reflect the correct structure of the environment. In addition to this, we also report the results of the probe sessions conducted in these experiments. Those results reveal how the pattern of choices is affected by the variations in the parameters.

The parameters used in these experiments are different from the parameters used in the main paper in two respects. Firstly, the inter-trial interval (ITI) in these supplementary experiments was non-zero, whereas the ITI in the experiment reported in the main paper was zero. Secondly, in the supplementary experiments auditory cues (clicker and tone) were used to signal stage 2 states, whereas in the experiments reported in the main paper visual cues were used to signal stage 2 states. See Behavioural procedures below for the details of the parameters used in each experiment.

In the supplementary experiments, because the ITI was non-zero, the stage 1 state (S0) was explicitly signalled to the animals at the end of the ITI by the illumination of the house light. We assumed that the first response made after the ITI was the state 1 action and the response after that was the stage 2 action (and this is what stage 1 actions and stage 2 actions refer to in analysis presented below). The data, however, indicated that animals did not wait for the presentation of the stage 1 state to make their first response and made a significant number of responses during the ITI. These were unrewarded regardless of which action/action sequence was made by the animals. Clearly, this could interfere with the formation and/or expression of action sequences and, therefore, in the experiment reported in the main paper an ITI of zero was used to remove such interference. Furthermore, we used visual cues in the experiment reported in the main paper instead of the auditory cues used in the supplementary experiments. This was because the visual cues (used in the experiment reported in the main paper) are presumably less salient and so harder to discriminate than the auditory cues used in the supplementary experiments; it took twice the number of training sessions for the animals to learn the discrimination between the visual cues compared to the auditory cues (comparing Figure A2 with Figure A3h). Because of this reduction in slaience, when visual cues were being used less interference was introduced during the transition between states and so there was a higher tendency for the rats to perform the stage 1 and stage 2 actions in a sequence (uninterrupted) independent of the stage 2 states, increasing the opportunity to detect action sequences. For this reason, in the experiment reported in the main paper we used visual cues.

##### A1.8.1 Subjects and apparatus

Details are similar to the experiment reported in the main paper (see section Subjects and apparatus).

##### A1.8.2 Behavioural procedures

The general structure of the experiments was similar to the experiment reported in the main paper and it is depicted in Figure 1. Below we explain each step and highlight the similarities and differences between the experiments.

Magazine training, lever-press training, and discrimination training phases (phases 1-3) were similar to the experiment reported in the main paper (see section Material and Methods), except that in the supplementary experiments, the stimuli were the tone and clicker, but in the experiment reported in the main paper, the stimuli were a constant and a blinking house light.

Next, the rats received training on the two-stage task in which animals first made a binary choice at stage 1 (signalled by the illumination of the house light), after which they transitioned to the stage 2 state, in which again they made another binary choice that could lead to either reward delivery or no-reward. In supplementary experiments 1 and 2 there was a 20-s inter-trial interval (ITI) before the next trial started, and there was a 5-second ITI in supplementary experiment 3. Stage 2 states were signalled by the stimuli trained in the previous phase of the experiment. The stage 2 states were presented immediately after stage 1 actions were taken, and the reward/no-reward was presented immediately after the stage 2 action was taken.

In each trial, only one of the stage 2 states led to reward, whereas the other state did not lead to reward irrespective of the choice of actions. The stage 2 states that earned a reward frequently switched between states during the course of a session (Figure 2a). In supplementary experiment 1, this switch occurred after every four outcomes with a maximum 40 outcomes in a session, which later in the training increased to every eight outcomes as shown in Figure A3a (with a maximum 48 outcomes in a session and maximum duration of a session was limited to an hour). In supplementary experiment 2, the switch occurred whenever a randomly selected number of outcomes were received since the last switch. This random number was uniformly drawn from within a range from 8 to 16 outcomes (maximum 48 outcomes in a session and maximum duration of a session was limited to an hour). In supplementary experiment 3, the switch occurred every fourth outcome received (maximum 50 outcomes in a session and maximum duration of a session was limited to an hour). Furthermore, because the ITI was long in supplementary experiments 1 and 2, animals received a pre-training phase on the two-stage task in which the reward in the stage 2 states was fixed during a session, and was changed across sessions. Subjects received ten training sessions in this manner. Similarly, in supplementary experiment 2, subjects received two pre-training sessions in which they could earn a reward in both stage 2 states.

Animals were trained on the two-stage task for 69 sessions in supplementary experiment 1, 57 sessions in supplementary experiment 2, 60 sessions in supplementary experiment 3. In the middle of, or at the end of these training sessions, animals were given probe test sessions in which stage 1 actions led to stage 2 states in a probabilistic manner. One stage 1 action led to its specific stage 2 state 80% of the time, whereas the other stage 1 action led to the other stage 2 state (Figure 2c). For the last probe session in Experiments 1 and 3, the probability that stage 1 actions led to the corresponding stage 2 states was 50%. The exact positions of probe sessions for supplementary experiments 1 to 3 are depicted in Figures A3b, A4b, A5b respectively marked with an asterisk.

##### A1.8.3 Results

Results for supplementary experiments 1-3 are shown in Figures A3, A4, A5 respectively. Multiple probe sessions were conducted in supplementary experiment 1 (marked with ‘*’ in Figure A3b) and the statistical analysis of probe sessions are shown in Table A5. Supplementary experiments 2,3 contained a single probe session (marked with * in Figure A4b, and Figure A5b), and the statistical analysis of these probe sessions are shown in Table A6. Note that the differences in the results of the probe sessions of the supplementary experiments compared to the those reported in the main paper is most likely due to the interference produced by unrewarded actions taken during the ITI (since the ITI was non-zero in the supplementary experiments), and also due to the use of auditory cues in the supplementary experiments (see above).

Similar to the experiment reported in main paper, at the beginning of training, as Figure A3a, A4a, A5a show, the rats not only failed to show a tendency to take the same action after earning a reward in the previous trial, they also showed a tendency to switch to the other action. This effect was statistically significant in the first five sessions of supplementary experiment 2 (sessions s20 to s24; *β* = *−*0.253 (CI: *−*0.371, *−*0.135), SE=0.060, p<10^*−*4^) and supplementary experiment 3 (sessions s15 to s19; *β* = *−*0.222 (CI: *−*0.357, *−*0.086), SE=0.068, p=0.001); but, for supplementary experiment 1, although the effects were in the same direction, it was not statistically significant (sessions s26 to s30; *β* = *−*0.063, SE=0.045, p=0.161), likely due to the pre-training the animals received in that experiment.

**Table A1.** For the best four models in each family, the table represents the negative log-model evidence (−log *P* (*D M*)) for each model, the number of free parameters of each model (df), the free parameters of each model, and the family of each model. The table also represents a pseudo-r statistic (p–*r*^2^), which is a normalized measure of the variance accounted for in comparison to a model with random choices (averaged over subjects). ‘*’ indicated the model with the best model evidence.

| model family | free-parameters | $p-r^2$ | df | $-\log P(D\vert M)$ |
| --- | --- | --- | --- | --- |
| HMB* | $\beta_1, \eta_1, k_2, \phi \ (I = 0)$ | 0.21 | 4 | 1172.09 |
| H-MB | $\beta_1, \eta_1, k_2, k_3, \phi \ (I = 0)$ | 0.21 | 5 | 1176.48 |
| HMB | $\beta_1, \eta_1, k_1, k_2, \phi \ (I = 0)$ | 0.21 | 5 | 1178.90 |
| H-MB | $\beta_1, \eta_1, k_2, \phi, I$ | 0.21 | 5 | 1178.93 |
| H | $\beta_1, \eta_1, k_2, \phi, I$ | 0.20 | 5 | 1185.27 |
| H | $\beta_1, \beta_2, \eta_1, k_2, I$ | 0.20 | 5 | 1187.93 |
| H | $\beta_1, \beta_2, \eta_1, k_1, \phi, I$ | 0.21 | 6 | 1190.48 |
| H | $\beta_1, \eta_1, \phi, I$ | 0.19 | 4 | 1195.04 |
| MF | $\beta_1, \beta_2, \alpha_1, \lambda$ | 0.17 | 4 | 1219.57 |
| MF | $\beta_1, \beta_2, \alpha_1, \lambda, k$ | 0.18 | 5 | 1220.41 |
| MB | $\beta_1, \beta_2, \eta, k$ | 0.18 | 4 | 1222.35 |
| MF-MB | $\beta_1, \beta_2, \alpha_1, \lambda, w$ | 0.18 | 5 | 1223.02 |
| MF-MB | $\beta_1, \beta_2, \alpha_1, \lambda, k, w$ | 0.19 | 6 | 1224.45 |
| MF | $\beta_1, \alpha_1, \lambda$ | 0.16 | 3 | 1224.62 |
| MF | $\beta_1, \alpha_1, \lambda, \phi$ | 0.17 | 4 | 1224.74 |
| MF-MB | $\beta_1, \alpha_1, \lambda, w$ | 0.17 | 4 | 1224.82 |
| MB | $\beta_1, \beta_2, \eta$ | 0.17 | 3 | 1225.02 |
| MB | $\beta_1, \eta, k, \phi$ | 0.17 | 4 | 1225.07 |
| MF-MB | $\beta_1, \alpha_1, \lambda, w, \phi$ | 0.18 | 5 | 1225.89 |
| MB | $\beta_1, \beta_2, \eta, \phi, k$ | 0.18 | 5 | 1226.82 |

**Table A2.** Value of the estimated parameters for each subject.

| subject | df | $p-r^2$ | no. choices | $-\log P(D_i\vert M, \Theta^{\text{ML}})$ | $\beta_1$ | $\eta_1$ | $\phi$ | $k_2$ |
| --- | --- | --- | --- | --- | --- | --- | --- | --- |
| 0 | 4 | 0.25 | 244 | 128.11 | 1.91 | 0.83 | 1.33 | 0.67 |
| 1 | 4 | 0.30 | 214 | 105.23 | 1.94 | 0.77 | 2.53 | 0.82 |
| 2 | 4 | 0.18 | 264 | 150.42 | 1.60 | 0.78 | 2.20 | 0.68 |
| 3 | 4 | 0.32 | 206 | 98.92 | 1.84 | 0.78 | 2.68 | 0.66 |
| 4 | 4 | 0.06 | 286 | 188.33 | 0.99 | 0.78 | 1.96 | 0.47 |
| 5 | 4 | 0.20 | 272 | 152.94 | 1.53 | 0.73 | 2.38 | 0.66 |
| 6 | 4 | 0.19 | 296 | 168.19 | 1.38 | 0.82 | 2.36 | 0.94 |
| 7 | 4 | 0.15 | 252 | 149.48 | 1.26 | 0.79 | 2.28 | 0.62 |

**Table A3.** Results of the logistic regression analysis of stage 1 and stage 2 choices for the experiment reported in the main paper. For the stage 1 choices, the analysis is focused on staying on the same stage 1 action on the next trial, based on whether the previous trial was rewarded (reward), and whether the previous trial was common or rare (transition). ‘reward:transition’ is the interaction between reward, and transition type, and ‘intercept’ refers to the intercept term. For stage 2 choices, the analysis is focused on staying on the same stage 2 action, based on staying on the same stage 1 action (stay) and earning a reward in the previous trial (reward). ‘reward:stay’ is the interaction between ‘reward’, and ‘stay’. ‘probe 3’ are the results reported in the main paper.

| Stage 1 actions |  |  |  |  |
| --- | --- | --- | --- | --- |
|  |  | probe 1 | probe 2 | probe 3 |
| intercept | *p*-value | 0.618 | 0.054 | 0.373 |
| | $\beta$ (SE) | -0.065 (0.131) | 0.299 (0.155) | -0.166 (0.186) |
| reward | *p*-value | 0.002 | <1e-4 | 0.003 |
| | $\beta$ (SE) | 0.52 (0.174) | 0.504 (0.128) | 0.612 (0.208) |
| transition | *p*-value | 0.832 | 0.155 | 0.164 |
| | $\beta$ (SE) | -0.057 (0.273) | -0.406 (0.285) | 0.371 (0.267) |
| reward:transition | *p*-value | 0.005 | 0.02 | <1e-5 |
| | $\beta$ (SE) | 0.842 (0.3) | 0.399 (0.171) | 0.923 (0.204) |
| Stage 2 actions |  |  |  |  |
|  |  | probe 1 | probe 2 | probe 3 |
| intercept | *p*-value | 0.221 | 0.576 | 0.001 |
| | $\beta$ (SE) | -0.266 (0.217) | -0.168 (0.301) | -0.95 (0.295) |
| reward | *p*-value | <1e-4 | <1e-5 | <0.001 |
| | $\beta$ (SE) | 0.631 (0.149) | 1.244 (0.265) | 0.849 (0.235) |
| stay | *p*-value | 0.111 | 0.255 | 0.019 |
| | $\beta$ (SE) | 0.254 (0.159) | 0.234 (0.205) | 0.608 (0.259) |
| reward:stay | *p*-value | 0.222 | 0.153 | 0.027 |
| | $\beta$ (SE) | 0.182 (0.149) | 0.29 (0.203) | 0.494 (0.224) |

**Figure A1.**
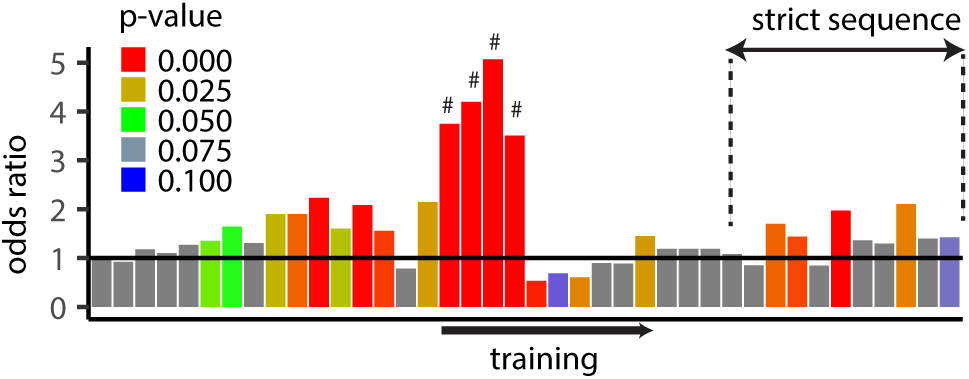
The graphs shows the intercept term for the analysis shown in Figure 3a, which is the odds ratio of staying on the same stage 1 action. The *p*-value of this analysis (for the intercept term) was used for colour-coding each bar, and the height of each bar represents the odds ratio calculated as *e*^*β*^, in which *β* is the coefficient obtained for the intercept term. The graphs can be interpreted as the probability of staying on the same stage 1 action, independent of whether reward was earn in the previous trial. The sessions marked with ‘#’ and ‘strict sequence’ are similar to the sessions described in Figure 3a.

**Table A4.** Results of the logistic regression analysis of stage 1 and stage 2 choices for the experiment reported in the main paper. For the stage 1 choices, the analysis is focused on staying on the same stage 1 action on the next trial, based on whether the previous trial was rewarded (reward), and whether the previous trial was common or rare (transition). ‘reward:transition’ is the interaction between reward, and transition type, and ‘intercept’ refers to the intercept term. ‘correct’ means that whether the correct stage 1 action was taken in the previous trial. ‘correct’ stage 1 action in refers to the stage 1 action which led the rewarded stage 2 state. For stage 2 choices, the analysis is focused on staying on the same stage 2 action, based on staying on the same stage 1 action (stay) and earning a reward in the previous trial (reward). ‘reward:stay’ is the interaction between ‘reward’, and ‘stay’. This table is different from Table A3 in two aspects: (i) the ‘correct’ predictor was added to the analysis following Akam et al. (2015) suggestion, (ii) unlike the analysis performed in Table A3, the trials in which subjects did not make the correct discrimination were not excluded from the analysis. Note that the aborted trial, i.e., the trials in which animals entered the magazine between stage 1 and stage 2 actions were excluded. The reason for this second difference is, since rewards are deterministic (within each reversal), if we only include trials in which animals make correct discrimination, then ‘reward-transition’ predictor, and ‘correct’ predictor will be identical. As such, in this analysis that ‘correct’ is a predictor, we included all the trials.

| Stage 1 actions |  |  |  |  |
| --- | --- | --- | --- | --- |
|  |  | probe 1 | probe 2 | probe 3 |
| intercept | p-value | 0.291 | 0.041 | 0.335 |
| | $\beta$ (SE) | -0.215 (0.204) | 0.304 (0.149) | -0.202 (0.209) |
| reward | p-value | <0.001 | <0.001 | 0.024 |
| | $\beta$ (SE) | 0.418 (0.115) | 0.447 (0.092) | 0.390 (0.174) |
| transition | p-value | 0.026 | 0.697 | 0.008 |
| | $\beta$ (SE) | 0.365 (0.164) | 0.055 (0.142) | 0.614 (0.235) |
| correct | p-value | 0.053 | 0.855 | 0.117 |
| | $\beta$ (SE) | 0.566 (0.293) | 0.048 (0.266) | 0.399 (0.255) |
| reward:transition | p-value | 0.625 | 0.777 | 0.026 |
| | $\beta$ (SE) | 0.113 (0.233) | -0.063 (0.225) | 0.527 (0.237) |
| Stage 2 actions |  |  |  |  |
|  |  | probe 1 | probe 2 | probe 3 |
| intercept | p-value | 0.156 | <0.001 | 0.296 |
| | $\beta$ (SE) | 0.153 (0.108) | 0.517 (0.138) | -0.189 (0.181) |
| reward | p-value | 0.144 | <0.001 | 0.315 |
| | $\beta$ (SE) | 0.172 (0.118) | 0.568 (0.126) | 0.125 (0.124) |
| stay | p-value | <0.001 | <0.001 | <0.001 |
| | $\beta$ (SE) | 0.390 (0.104) | 0.406 (0.110) | 0.673 (0.187) |
| reward:stay | p-value | 0.361 | 0.264 | <0.001 |
| | $\beta$ (SE) | 0.096 (0.105) | 0.131 (0.118) | 0.408 (0.121) |

**Table A5.** Results of the logistic regression analysis of stage 1, and stage 2 choices in supplementary experiment For the stage 1 choices, the analysis is focused on staying on the same stage 1 action on the next trial based on whether the previous trial was rewarded (reward) and whether the previous trial was common or rare (transition). ‘reward:transition’ is the interaction between reward, and transition type. For stage 2 choices, the analysis focuses on staying on the same stage 2 action, based on staying on the same stage 1 action (stay), and earning a reward on the previous trial (reward). ‘reward:stay’ is the interaction between ‘reward’, and ‘stay’. ‘*p*’ refers to *p*-value. For the last probe session (s94) since the pattern of choices was not stationary but changing during the session, we presented separately the analysis for the first 16 earned outcomes during the sessions (s94:1; outcomes 1:16), second 16 outcomes earned during the session (s94:2; outcomes 17:32), and third 16 outcomes earned during the session (s94:3; outcomes 33:48).

| Stage 1 actions |  |  |  |  |  |  |  |  |  |  |  |  |
| --- | --- | --- | --- | --- | --- | --- | --- | --- | --- | --- | --- | --- |
| session |  | s32 | S40 | s49 | s57 | s66 | s78 | s87 | s94 | s94:1 | s94:2 | s94:3 |
| intercept | $p$ | 0.773 | 0.312 | 0.017 | 0.299 | 0.177 | 0.162 | 0.885 | 0.428 | 0.46 | 0.131 | 0.194 |
|  | SE | 0.151 | 0.11 | 0.16 | 0.124 | 0.136 | 0.11 | 0.228 | 0.127 | 0.248 | 0.263 | 0.29 |
| | $\beta$ | 0.043 | 0.111 | 0.382 | 0.129 | 0.184 | 0.15 | -0.032 | 0.1 | 0.183 | -0.397 | 0.377 |
| reward | $p$ | 0.938 | 0.085 | 0.09 | 0.067 | 0.021 | 0.001 | 0.904 | <0.001 | 0.356 | 0.003 | 0.007 |
|  | SE | 0.133 | 0.117 | 0.133 | 0.145 | 0.117 | 0.12 | 0.204 | 0.121 | 0.207 | 0.295 | 0.234 |
| | $\beta$ | -0.01 | 0.202 | 0.226 | 0.265 | 0.27 | 0.39 | 0.024 | 0.465 | 0.191 | 0.854 | 0.627 |
| transition | $p$ | 0.764 | 0.628 | 0.098 | 0.016 | 0.825 | 0.383 | 0.993 | 0.498 | 0.407 | 0.644 | 0.335 |
|  | SE | 0.14 | 0.105 | 0.191 | 0.119 | 0.124 | 0.13 | 0.24 | 0.14 | 0.187 | 0.296 | 0.185 |
| | $\beta$ | 0.042 | -0.051 | -0.315 | -0.285 | -0.027 | -0.11 | 0.002 | 0.094 | 0.155 | -0.136 | 0.178 |
| reward:transition | $p$ | 0.882 | 0.013 | 0.362 | 0.397 | 0.485 | 0.417 | <0.001 | <0.001 | 0.011 | 0.003 | 0.668 |
|  | SE | 0.129 | 0.113 | 0.141 | 0.129 | 0.107 | 0.09 | 0.182 | 0.114 | 0.181 | 0.29 | 0.212 |
| | $\beta$ | 0.019 | -0.281 | -0.129 | 0.109 | -0.075 | 0.08 | 0.623 | 0.404 | 0.459 | 0.85 | 0.091 |
| Stage 2 actions |  |  |  |  |  |  |  |  |  |  |  |  |
| session |  | s32 | s40 | s49 | s57 | s66 | s78 | s87 | s94 | s94:1 | s94:2 | s94:3 |
| intercept | $p$ | <1e-7 | <1e-12 | <1e-11 | 0.963 | <1e-8 | <1e-14 | <1e-9 | <1e-16 | 0.969 | <1e-4 | 0.972 |
|  | SE | 0.446 | 0.411 | 0.434 | 190.44 | 0.477 | 0.35 | 0.459 | 0.293 | 195.286 | 0.577 | 514.163 |
| | $\beta$ | -2.449 | -2.936 | -2.996 | -8.943 | -2.848 | -2.82 | -2.969 | -2.672 | -7.546 | -2.477 | -17.949 |
| reward | $p$ | 0.933 | 0.938 | 0.148 | 0.984 | 0.383 | 0.137 | 0.169 | 0.019 | 0.98 | 0.924 | 0.995 |
|  | SE | 0.327 | 0.379 | 0.459 | 190.461 | 0.433 | 0.32 | 0.459 | 0.268 | 195.285 | 0.512 | 740.305 |
| | $\beta$ | 0.027 | 0.029 | 0.662 | 3.825 | 0.377 | 0.48 | 0.631 | 0.626 | 5.015 | 0.049 | 4.749 |
| stay | $p$ | 0.057 | 0.852 | 0.074 | 0.973 | 0.401 | 0.386 | 0.891 | 0.798 | 0.98 | 0.387 | 0.994 |
|  | SE | 0.406 | 0.522 | 0.487 | 190.44 | 0.405 | 0.33 | 0.463 | 0.278 | 195.286 | 0.507 | 665.334 |
| | $\beta$ | 0.771 | -0.097 | 0.868 | -6.429 | 0.339 | 0.29 | 0.063 | 0.071 | -4.891 | 0.438 | 5.35 |
| reward:stay | $p$ | 0.105 | 0.515 | 0.957 | 0.982 | 0.089 | 0.2 | 0.443 | 0.853 | 0.984 | 0.809 | 0.994 |
|  | SE | 0.32 | 0.356 | 0.454 | 190.46 | 0.398 | 0.37 | 0.49 | 0.283 | 195.286 | 0.503 | 615.096 |
| | $\beta$ | 0.519 | 0.232 | -0.024 | 4.252 | 0.677 | 0.47 | 0.376 | -0.052 | 3.974 | -0.121 | 4.881 |

**Table A6.**
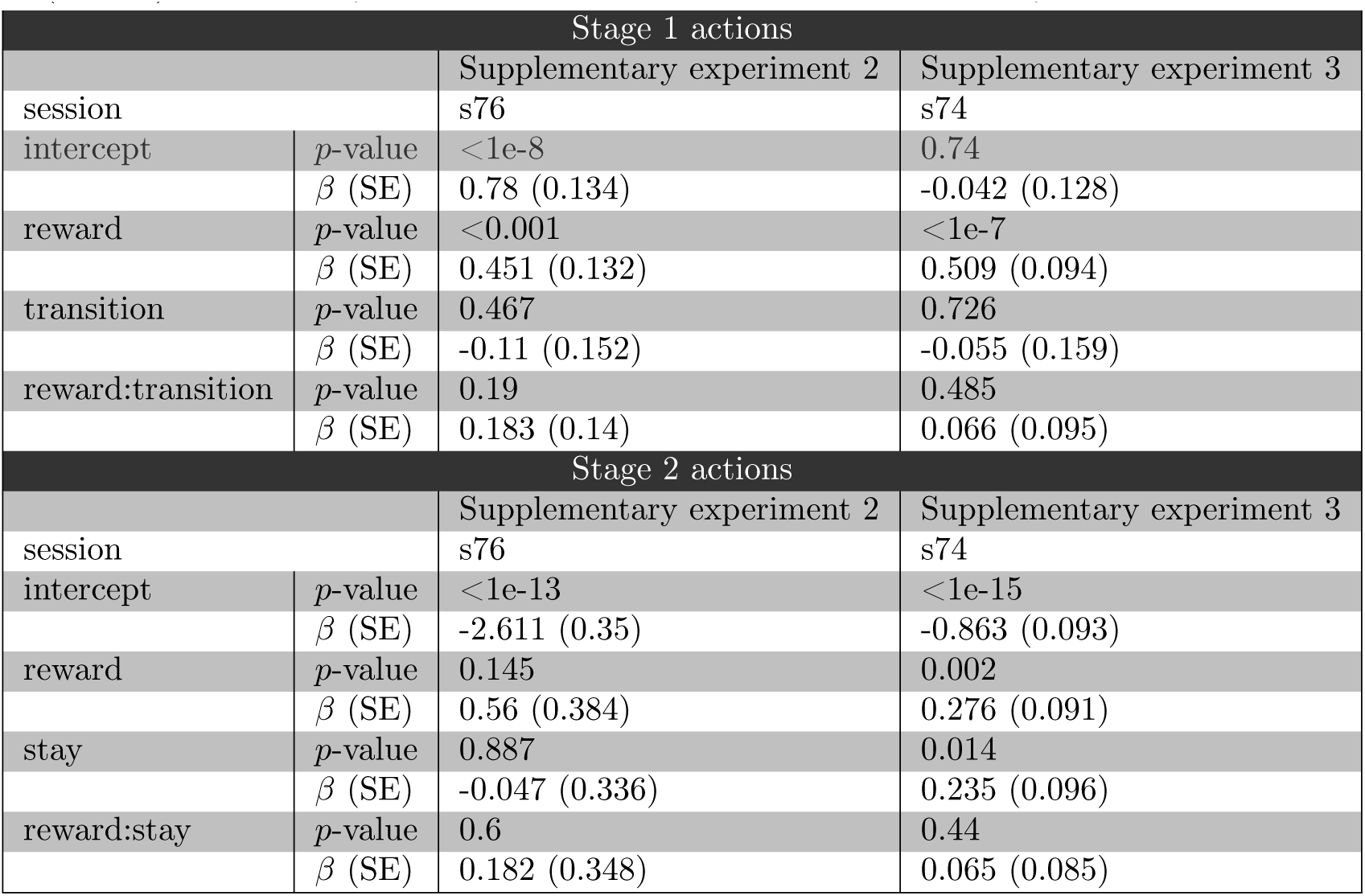
Results of the logistic regression analysis of stage 1 and stage 2 choices in supplementary experiments 2 and 3. For the stage 1 choices, the analysis is focused on staying on the same stage 1 action on the next trial, based on whether the previous trial was rewarded (reward), and whether the previous trial was common or rare (transition). ‘reward:transition’ is the interaction between reward, and transition type. For stage 2 choices, the analysis is focused on staying on the same stage 2 action, based on staying on the same stage 1 action (stay) and earning a reward in the previous trial (reward). ‘reward:stay’ is the interaction between ‘reward’, and ‘stay’.

**Figure A2.**
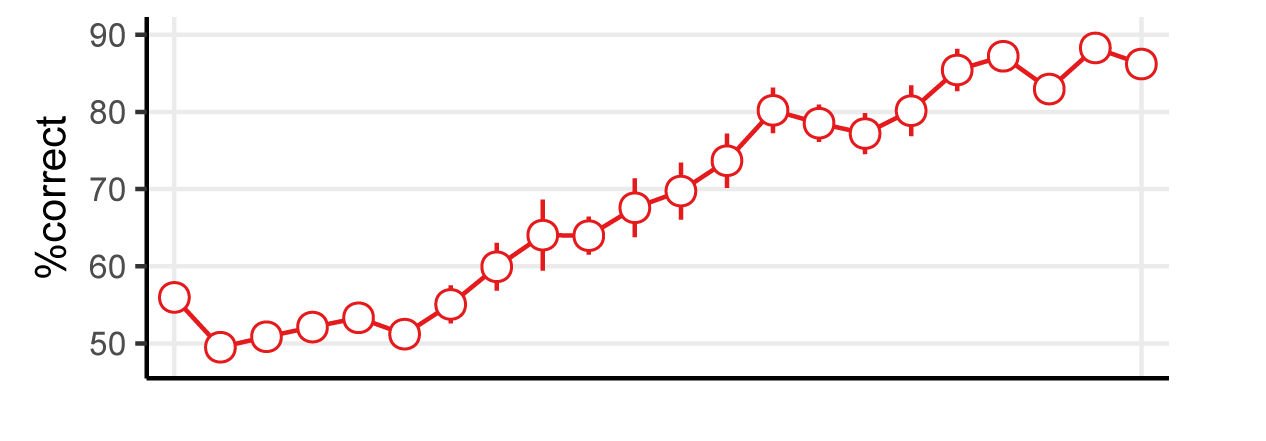
Results of discrimination training (for the experiment reported in the main paper) showing the percentage of correct responses averaged over subjects. Each point refers to a training session and error-bars are *±*1 SEM.

**Figure A3.**
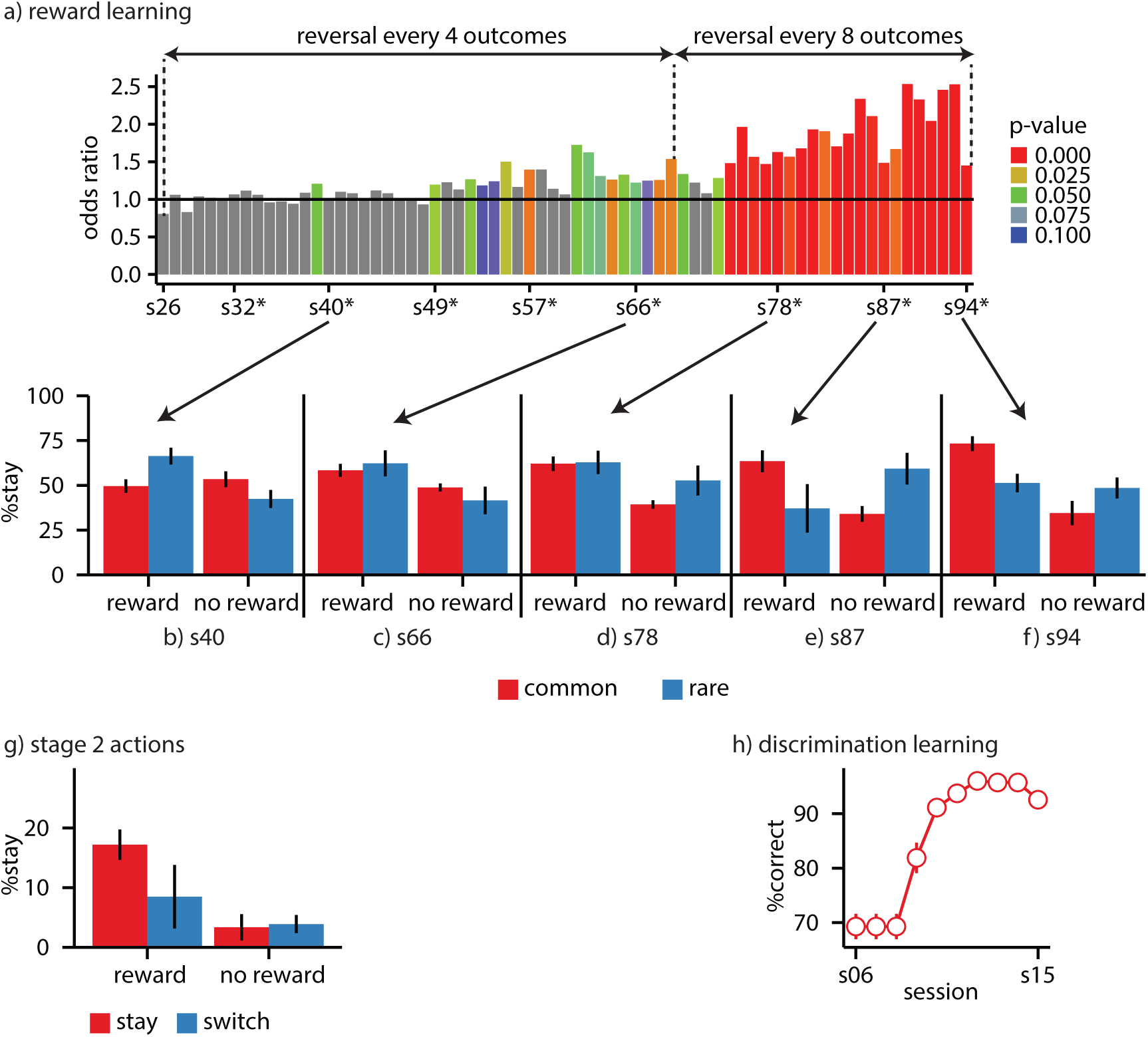
Supplementary experiment 1. In this experiment several probe sessions were inserted in the middle of the training sessions in order illustrate the development of actions over the training period. (a) Odds ratio of the probability of staying on the same stage 1 action after getting rewarded on the previous trial. Odds ratio=1 implies an equal preference for both actions. Sessions marked with ‘*’ are the probe sessions which included both rare and common transitions. (b-f) The probability of staying on the same stage 1 action in the probe sessions, averaged over subjects, as a function of whether the previous trial was rewarded (reward/no reward), and whether the transition in the previous trial was common or rare. The graphs illustrate a gradual shift from a simple state-space representation (panel b) to reward-guided actions (panels c,d), to goal-directed choices (panel e), and finally to a mixture of goal-directed and automatic actions (panel f). (g) Stage 2 actions in session s78. The graph shows the probability of staying on the same stage 2 action, averaged over subjects, as a function of whether the previous trial was rewarded (reward/no reward), and whether subjects stayed on the same stage 1 action (stay/switch). Similar to the analysis of the experiment reported in the main paper, only trials in which the stage 2 state is different from the previous trial are included in panel (g) in order to detect the performance of action sequences. Similarly, only trials in which subjects made a correct discrimination on the previous trial (‘R’ in S2, and ‘L’ in S1) were included in panels (a-g; see text). In all the probe sessions, the probability of rare transitions was 80%, except for the last session (s94) in which the probability of common and rare transitions was equal (i.e., 50%), in order to establish the effect of the transition probabilities on actions. (h) Results of initial discrimination training showing the percentage of correct responses averaged over subjects. Error-bars *±*1 SEM.

**Figure A4.**
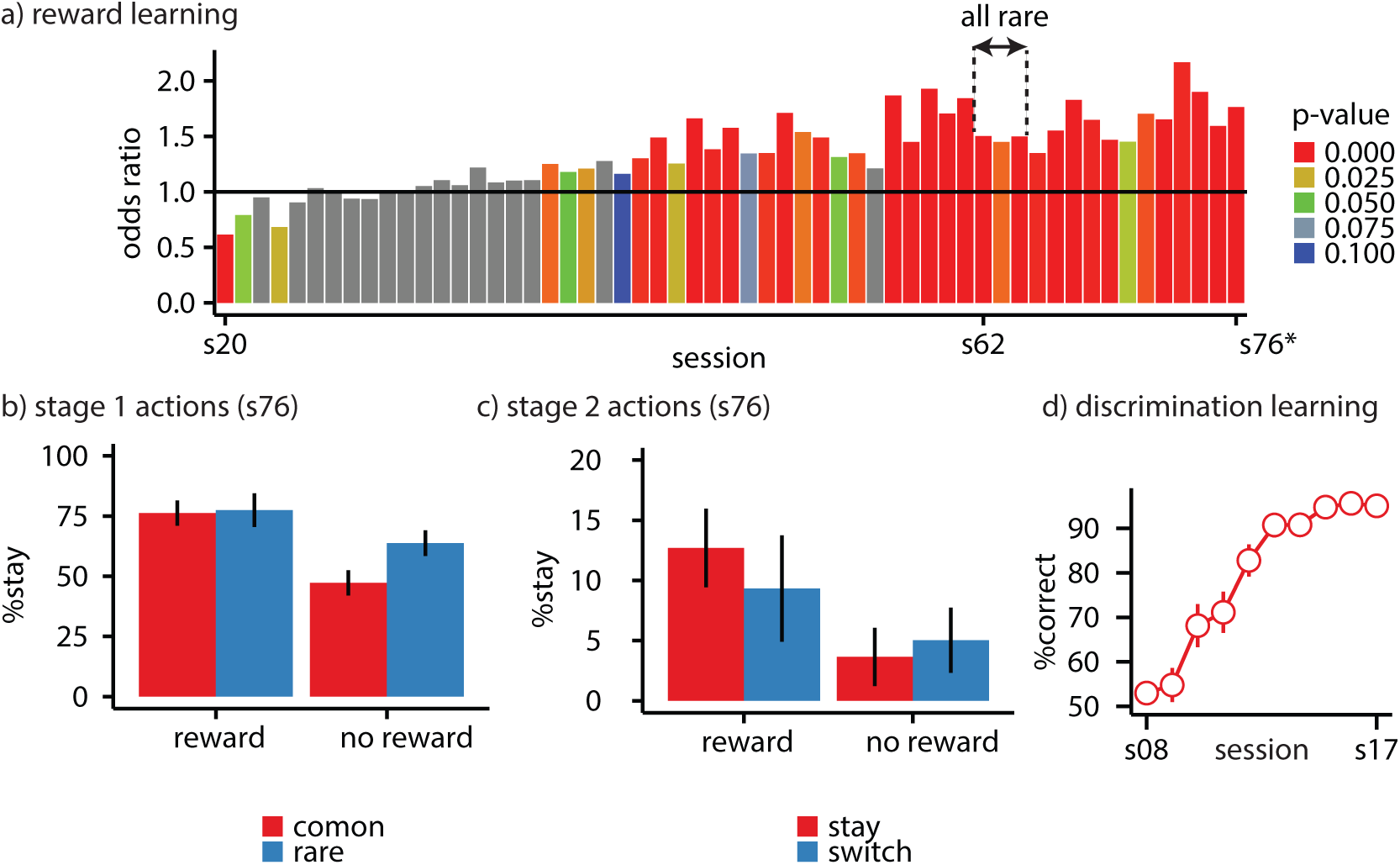
Supplementary experiment 2. (a) The odds ratio of staying on the same stage 1 action after earning a reward on the previous trial over the odds after earning no reward. Sessions denoted by ‘all rare’ included only rare transitions (similar to sessions marked with ‘#’ in Figure 3a). (b) The probability of staying on the same stage 1 action in the probe session (session s76) as a function of whether the previous trial was rewarded (reward/no reward) and whether the transition in the previous trial was common or rare. (c) The probability of staying on the same stage 2 action in the probe session (session s76), as a function of whether the previous trial was rewarded (reward/no reward) and whether subjects stayed on the same stage 1 action (stay/switch). Similar to the analysis in the main paper, only trials in which the stage 2 state was different from the previous trial are included in panels in order to detect the performance of action sequences. Similarly, only trials in which subjects made a correct discrimination on the previous trial (‘R’ in S2, and ‘L’ in S1) were included in panels (a-c). In all the probe sessions the probability of rare transitions was 50%. (d) Results of discrimination training showing the percentage of correct responses. Error-bars *±*1 SEM.

**Figure A5.**
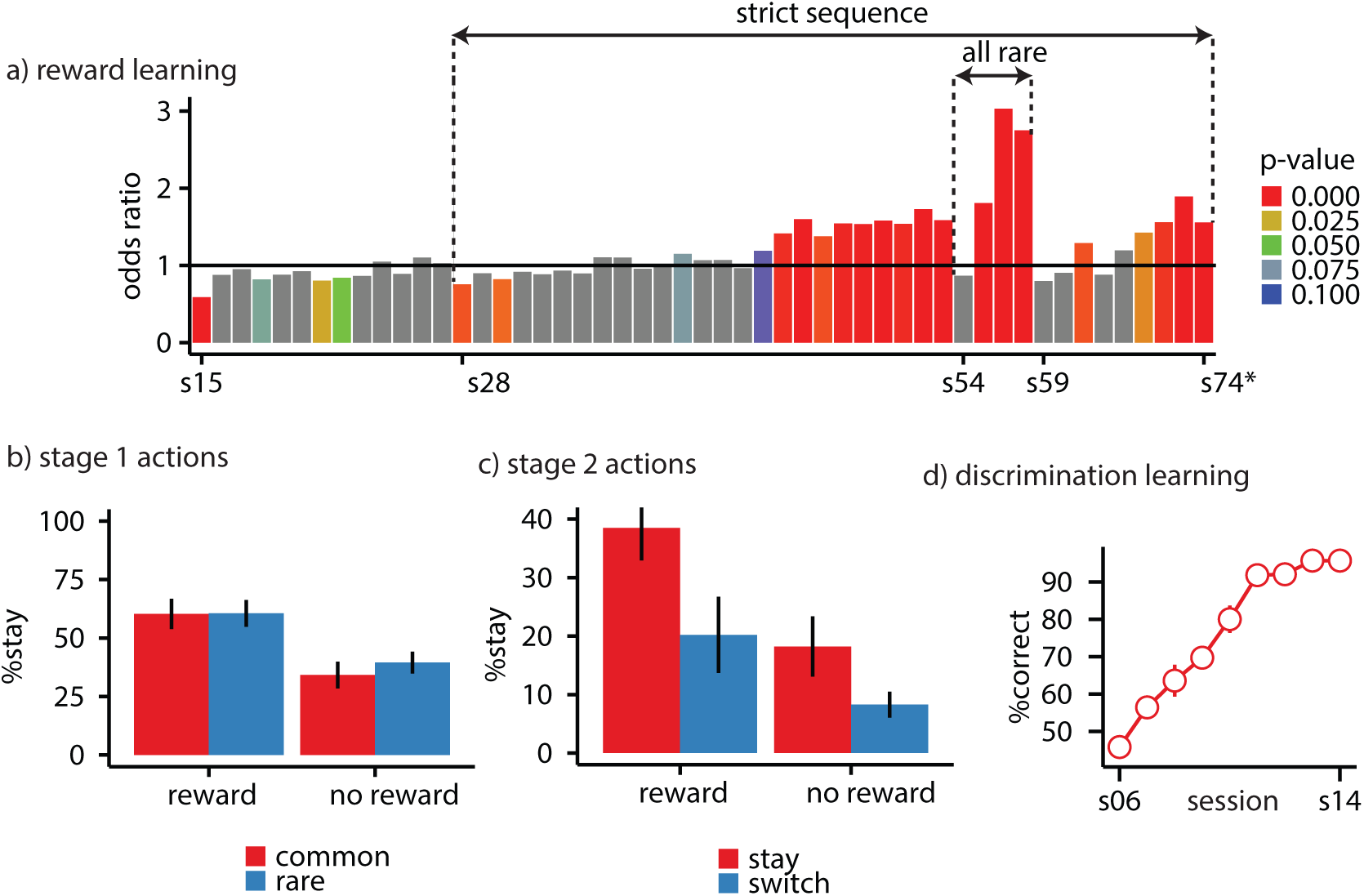
Supplementary experiment 3. (a) The odds ratio of staying on the same stage 1 action after earning a reward on the previous trial over the odds after earning no reward. Sessions denoted by ‘all rare’ included only rare transitions (similar to sessions marked with ‘#’ in Figure 3a). ‘strict sequences’ indicates that trials with magazine responses after stage 1 actions were aborted. (b) The probability of staying on the same stage 1 action in the probe session (session s74) as a function of whether the previous trial was rewarded (reward/no reward), and whether the transition in the previous trial was common or rare. (c) The probability of staying on the same stage 2 action in the probe session (session s74), as a function of whether the previous trial was rewarded (reward/no reward), and whether subjects stayed on the same stage 1 action (stay/switch). Similar to the analysis presented in the main paper, only trials in which the stage 2 states were different from the previous trial are included in panels (c) in order to detect the performance of action sequences. Similar to the analysis in the main paper, only trials in which subjects made a correct discrimination on the previous trial (‘R’ in S2, and ‘L’ in S1) were included in panels (a-c). In the probe sessions, the probability of rare transitions was 50%. (d) Results of discrimination training showing the percentage of correct responses. Error-bars *±*1 SEM.

